# Ion-channel transcript loss and variable firing in SCA1 Purkinje cells: a modelling test

**DOI:** 10.64898/2026.11.22.768239

**Authors:** Omkar Chandra, Caleb M. Smith

## Abstract

In spinocerebellar ataxia type 1 (SCA1), Purkinje cells fire with more variable interspike intervals before they degenerate, and reduced ion-channel expression is a proposed cause. We asked whether their channel transcript changes, applied as conductance changes, could produce this firing. We analysed public single-nucleus RNA-seq from male and female Atxn1 154Q/2Q knock-in and wild-type mice (24 discovery mice, 5–30 weeks; six held-out, 12 weeks). Knock-in Purkinje nuclei showed a reproducible transcriptional shift from 5 weeks. Transcripts of five channels, Kv3.3, BK, Kv4.3, Cav2.1 and Cav3.1, were reduced by 12 weeks (28–41 % at 18 weeks) and also lower, though not significantly for any single transcript, in ATXN1[82Q] Purkinje nuclei. We scaled the five conductances by these ratios in a published Purkinje cell model (130 parameter sets). The simulated neurons fired at 1.9 times the wild-type rate, outside the range recorded in knock-in Purkinje cells, and at 1.14 times in a second model. Scaling only Kv3.3, BK, Kv4.3 and Cav3.1, with Cav2.1 left at its wild-type value, brought both models close to the wild-type rate. With BK and Cav3.1 set from published staining, the afterhyperpolarization and depolarization-block threshold moved towards knock-in values, but by less. Dividing the five transcript ratios by the smaller knock-in membrane area (0.80 of wild type) also brought the median firing rate into the recorded range. Among versions whose median firing rate was in the recorded range, the median over parameter sets of the KI/WT ratio of the interspike-interval coefficient of variation was at most 1.57, whereas the recorded ratios of the group means were 1.8 and 2.9. In some versions, pauses in a few simulated neurons raised the ratio of the mean coefficients of variation. In these models, transcript changes alone did not reproduce the firing. Measuring spontaneous firing, Cav2.1 current density and membrane capacitance in the same knock-in and wild-type Purkinje cells, matched for age and lobule, would show whether Cav2.1 current falls less than its transcript, whether the cells are smaller, or both.

**Significance Statement:** In spinocerebellar ataxia type 1 (SCA1), Purkinje cells fire with more variable interspike intervals before they degenerate. Five ion-channel transcripts are lower in SCA1 knock-in Purkinje cells and, though not significantly for any single transcript, in a second SCA1 model. We asked whether these lower transcripts, taken as smaller conductances, explain the firing. In a Purkinje cell model, reducing all five conductances made the simulated neurons fire almost twice as fast as wild type, unlike knock-in Purkinje cells. Keeping Cav2.1 at its wild-type value, or correcting for the smaller knock-in membrane, brought the median firing rate close to that of knock-in cells, but the median increase in the variability of the interspike interval stayed below the recorded increase. Measuring spontaneous firing, Cav2.1 current density and membrane capacitance in the same knock-in and wild-type Purkinje cells, matched for age and lobule, would show whether Cav2.1 current falls less than its transcript, whether the cells are smaller, or both.

## Introduction

The hereditary disease spinocerebellar ataxia type 1 (SCA1) results from an expanded polyglutamine stretch in ATXN1, which causes progressive degeneration of cerebellar Purkinje cells. Changes in Purkinje cell firing before cell loss differ across mouse models (Hourez et al. 2011; Dell’Orco et al. 2015; Bushart et al. 2021). In the Atxn1 154Q/2Q knock-in (Watase et al. 2002), Purkinje cells at 14 weeks fire at a rate similar to that of wild-type cells but with more variable interspike intervals and a more depolarized afterhyperpolarization, and they reach depolarization block with a smaller injected current (Bushart et al. 2021). Staining in the same study showed decreased BK and Cav3.1 in the molecular layer but no significant change in Cav2.1 or IP3 receptor. In the ATXN1[82Q] transgenic line, many Purkinje cells are silent or in depolarization block at the onset of disease and later partially recover firing as they atrophy (Dell’Orco et al. 2015; Chopra et al. 2020). Across ataxias, reduced function of two classes of calcium-activated potassium channels, the large-conductance BK channel (Kcnma1) and the small-conductance (SK) channels, has been linked to these firing changes (Walter et al. 2006; Shakkottai et al. 2011; Dell’Orco et al. 2015; Bushart et al. 2021). Ion-channel dysfunction and altered Purkinje cell firing in ataxias have been reviewed (Hoxha et al. 2018; Bushart and Shakkottai 2019; Cook et al. 2021).

Single-nucleus RNA-seq (snRNA-seq) gives the change in ion-channel transcripts per cell type and age, but a transcript is not a conductance. mRNA levels explain only about half of the variation in protein levels across genes (Schwanhäusser et al. 2011; Liu et al. 2016), and the final conductance of a cell also depends on translation, trafficking, auxiliary subunits and membrane area. Channel density and transcript level diverge with decreasing size of the dendrite (Dell’Orco et al. 2015). If the step from RNA to conductance is stated as an assumption, a conductance-based model can test whether a transcript signature is enough to produce the reported phenotype. Disease mRNA ratios have been tested in populations of cardiac cell models (Walmsley et al. 2013), and various protein reductions have been modelled for SCA1 calcium signalling (Brown and Loew 2012), but we found no study that has applied transcript ratios measured for each gene and age to a population of Purkinje cell models and then restored each channel individually.

Ion-channel gene dysregulation has been reported across SCA1 and SCA2 mouse models (Chopra et al. 2020). Using the snRNA-seq time course of Tejwani et al. (2024), who described the transcriptional changes of each cerebellar cell type during disease, we compared the transcripts of Purkinje cells of Atxn1 154Q/2Q knock-in (KI) and wild-type (WT) mice, with the mouse as the unit and with replication in held-out mice. We asked whether these changes, applied as conductance changes in a published Purkinje cell model, could reproduce the firing rate, the coefficient of variation (CV) of the interspike interval, the afterhyperpolarization and the threshold to depolarization block recorded in KI Purkinje cells of the same line (Bushart et al. 2021). We then restored the conductances singly and in combination to find which reductions are compatible with the recordings.

## Results

### A reproducible genotype-associated transcriptional shift in Purkinje nuclei

To assess whether the RNA-level changes, used to scale a simulated Purkinje neuron, were sufficient to recreate the pathological firing phenotype recorded electrophysiologically in Purkinje cells of the same knock-in line (Bushart et al. 2021), we used publicly available, processed snRNA-seq data from SCA1 mouse cerebellum (GEO GSE246183; Tejwani et al. 2024). Figure 1 shows our study design and analyses.

**Figure 1.**
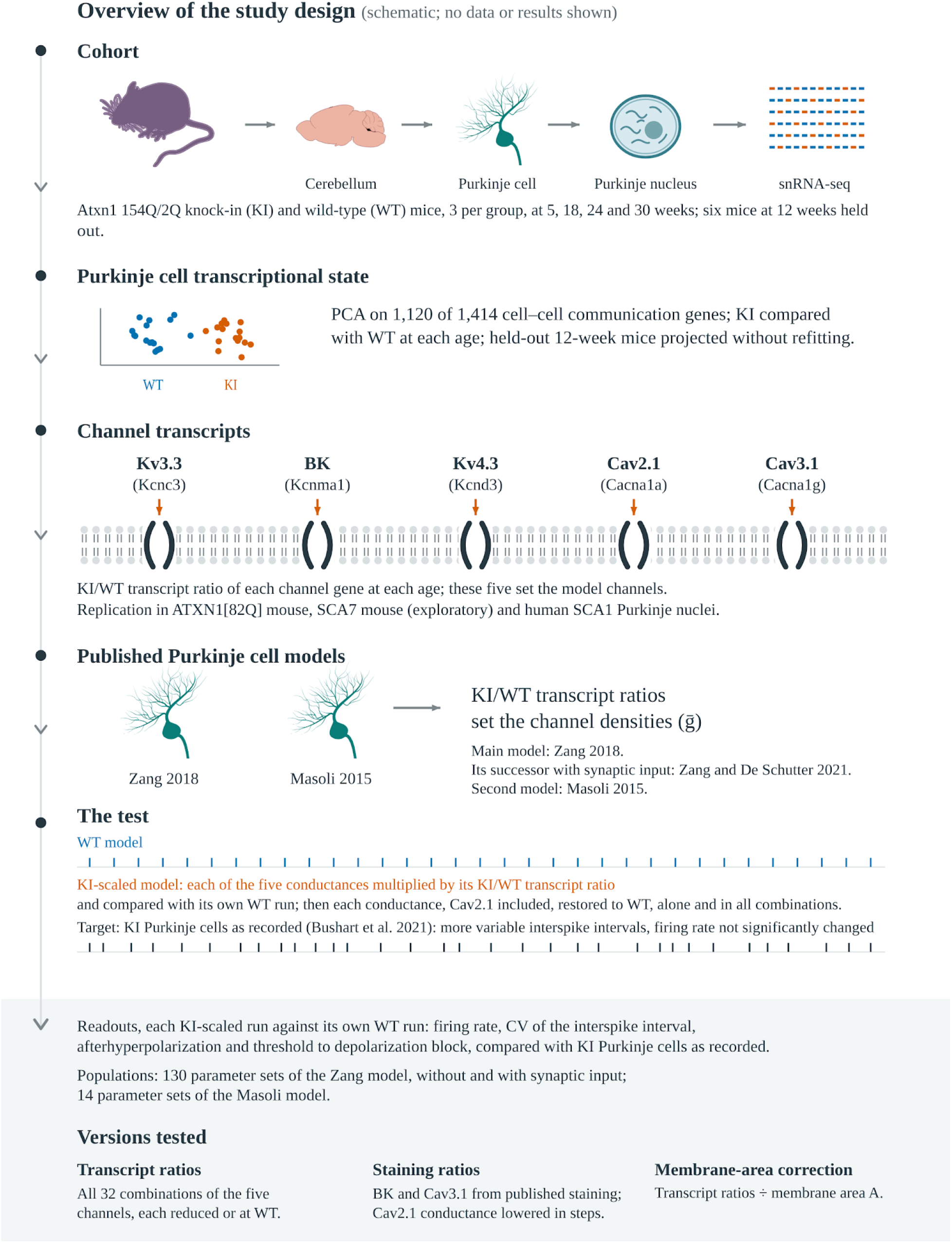
Overview of the study design. The figure shows what was done, not the results. Cohort: Atxn1 154Q/2Q knock-in (KI) and wild-type (WT) mice (three per genotype) at 5, 18, 24 and 30 weeks, and six 12-week mice held out of the discovery analyses; snRNA-seq of cerebellar Purkinje nuclei. Transcriptional state: principal component analysis of the 1,120 expressed genes of a panel of 1,414 cell–cell communication genes, with the held-out mice projected without refitting. Channel transcripts: the KI/WT transcript ratios of the five channel genes set the channels of the models. Replication was tested in ATXN1[82Q], SCA7 (exploratory) and human SCA1 Purkinje nuclei. Models: the main model of Zang et al. (2018) and its successor with synaptic input (Zang and De Schutter 2021), and the model of Masoli et al. (2015; Methods). The test: each KI-scaled run, with the five conductances multiplied by their transcript ratios, was compared with its own WT run. Each conductance was then restored to its WT value, alone and in all combinations, including Cav2.1 at its WT value with the other four reduced. Target: KI Purkinje cells as recorded (Bushart et al. 2021), with more variable interspike intervals than WT cells at a firing rate not significantly different from WT. Readouts: firing rate, CV of the interspike interval, afterhyperpolarization and threshold to depolarization block. Populations: 130 parameter sets of the Zang model, without and with synaptic input, and 14 of the Masoli model. Versions tested: all 32 combinations of the five transcript ratios; BK and Cav3.1 at their staining ratios, with Cav2.1 lowered in steps; the five transcript ratios divided by the membrane area A. Spike rasters and drawings are schematic. Mouse silhouette from NIH BioArt (public domain); Purkinje cell drawing adapted from A. Markanday, SciDraw (CC BY 4.0, doi:10.5281/zenodo.3926003).

Before scaling the simulated neurons by the transcript ratios, we asked whether KI Purkinje cells occupy a distinct transcriptional state (Figure 2). We performed principal component analysis of the expression of the 1,120 expressed genes of a panel of 1,414 genes involved in cell–cell communication (genes encoding ligands, receptors, ion channels, pumps and transporters, and synaptic proteins, listed from public databases before KI was compared with WT; Methods) in the Purkinje nuclei of the 24 discovery mice (Table 1). Every comparison was made between mice (three per genotype and age), never between nuclei, and the six 12-week mice were excluded from these fits. Of the 30 principal components kept, we called a component a genotype axis if the per-mouse means of the three KI and the three WT mice separated completely, in the same direction, at 3 or more of the 4 discovery ages.

**Figure 2.**
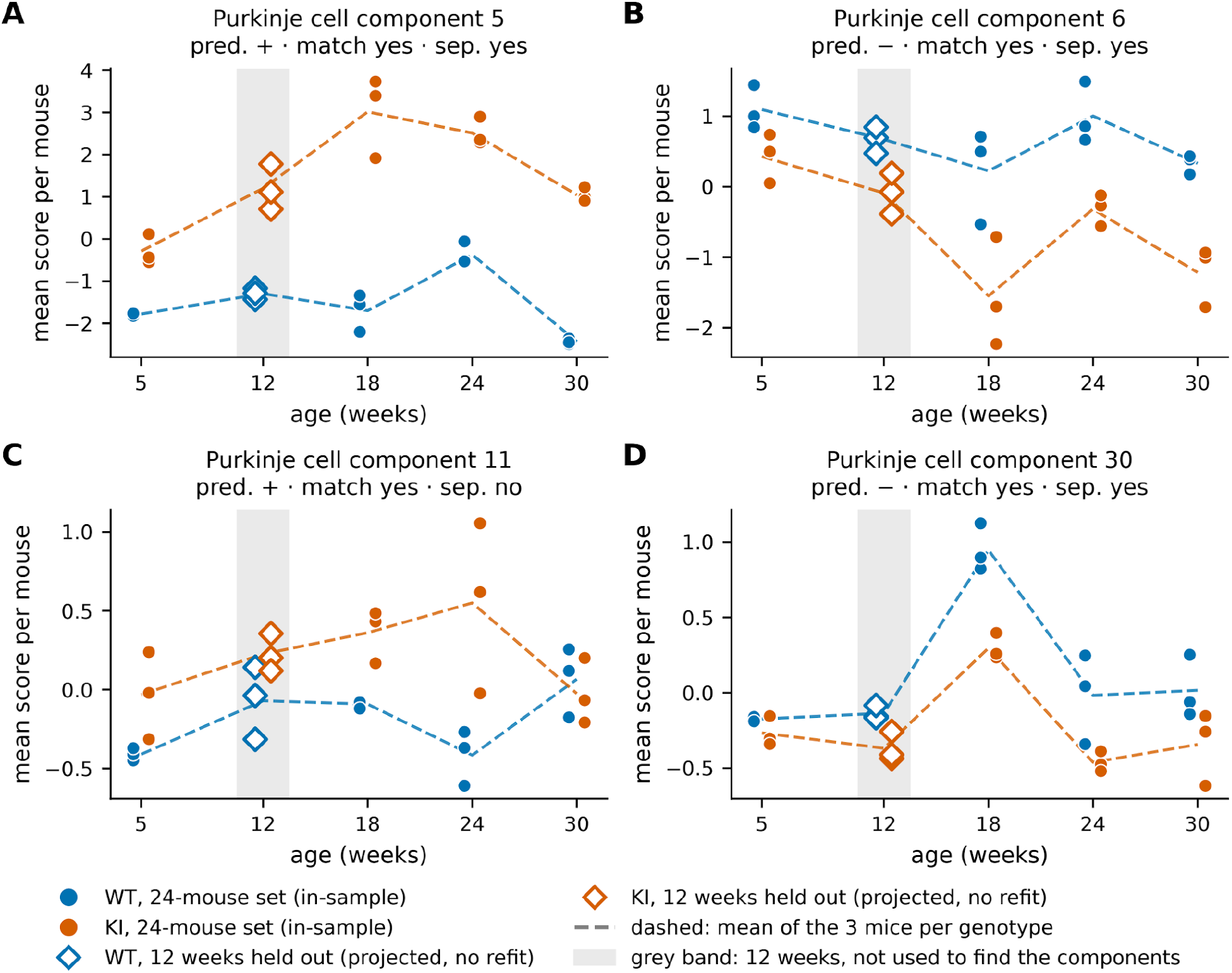
Genotype components in Purkinje nuclei. (A–D) Mean component score per mouse against age for components 5, 6, 11 and 30. Filled circles are individual discovery mice (n = 3 per genotype and age; blue, WT; orange, KI), and dashed lines join the means of the three mice of each genotype. Open diamonds in the grey band are the six held-out 12-week mice, which were projected onto the same components without refitting and were not used to find them. In the panel captions, “pred.” is the predicted sign of KI − WT (the majority sign over 5, 18, 24 and 30 weeks), “match” means agreement of the 12-week sign with the prediction, and “sep.” means complete 3-versus-3 separation of the 12-week mice in the predicted direction. See Extended Data Figure 2-1 for the full table, Extended Data Figure 2-2 for the gene loadings and Extended Data Figure 2-3 for the distribution of the nuclei of each mouse along the axis direction.

**Table 1.** Cohort and data.

| Item | Value |
|---|---|
| Source | GEO GSE246183: Atxn1 154Q/2Q knock-in and WT mouse cerebellum (Tejwani et al. 2024); the authors’ processed object, used from its raw UMI counts with the authors’ cell-type labels |
| Discovery mice | 24: 3 KI and 3 WT at each of 5, 18, 24 and 30 weeks |
| Held-out mice | 6: 3 KI and 3 WT at 12 weeks, held out of the discovery analyses |
| Nuclei after quality control | 280,295 discovery nuclei in 13 cell types; about 38,000 at 12 weeks |
| Focus population | Purkinje cells (1,971 discovery; 2,174 with 12 weeks) |
| Unit of inference | The mouse; nuclei are never treated as replicates |
| Known confounds | Sex confounded with age (18 weeks male, 30 weeks female); one library per mouse; ambient RNA not estimable |

Four components met this rule in Purkinje cells (Figure 2). Projected onto these components without refitting, the six held-out 12-week mice fell in the predicted direction on all 4 components, and the three KI mice separated completely from the three WT mice on 3 of them (Figure 2; Extended Data Figure 2-1 gives the counts expected by chance). One component, component 5, separated KI from WT at every age. Among its genes with the largest negative loadings are BK β4 (Kcnmb4) and IP3 receptor 1 (Itpr1), which are both reduced in KI Purkinje cells (Extended Data Figures 2-2 and 3-1). The four components share many genes, so these counts are not independent, and we report them as effect sizes, not as one p-value.

**Figure 3.**
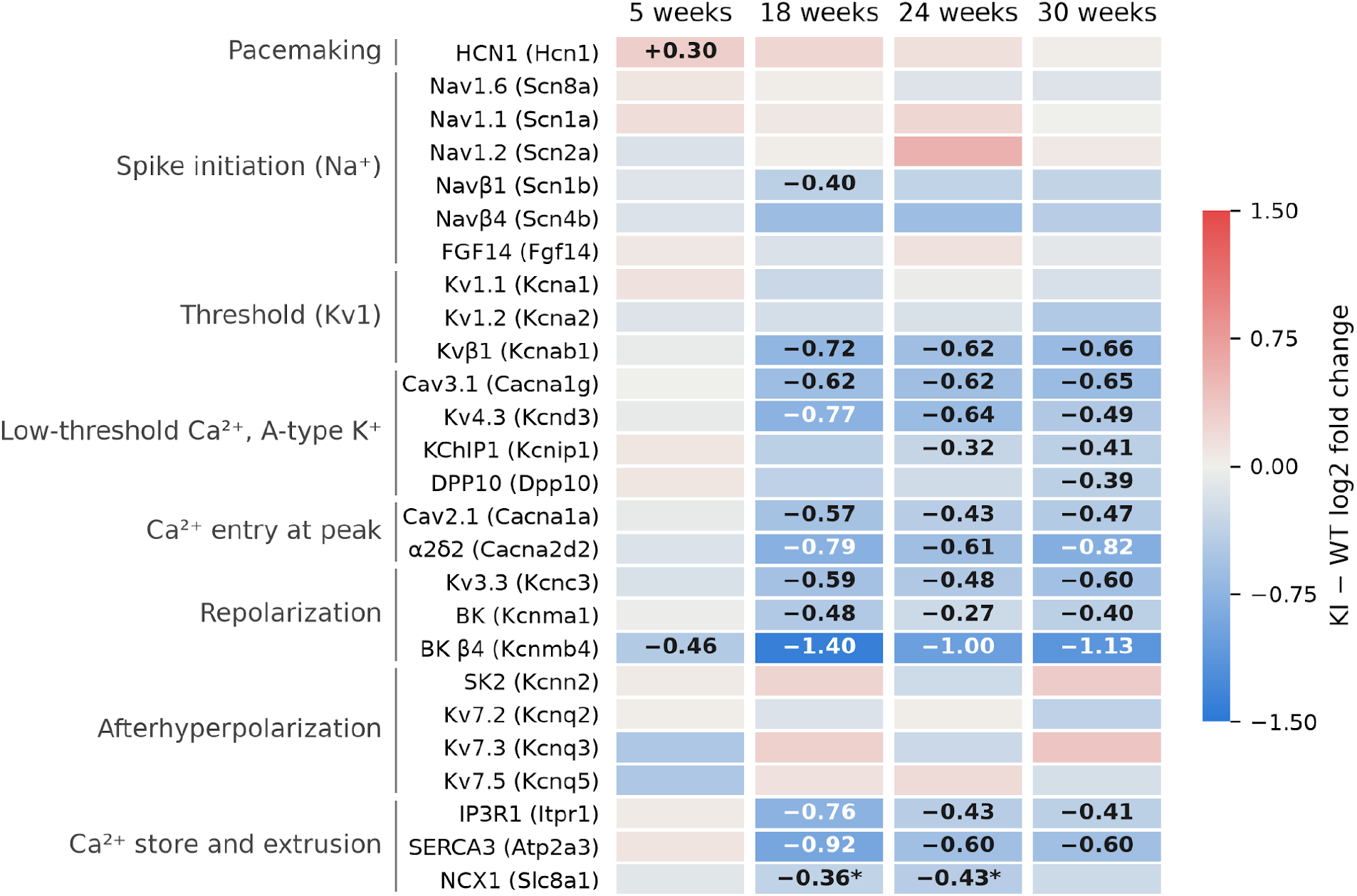
Ion-channel and calcium-store transcripts in KI Purkinje nuclei. KI − WT log2 fold change of 26 transcripts in Purkinje nuclei at 5, 18, 24 and 30 weeks. Rows are labelled with the channel or protein and, in parentheses, its gene. Rows are grouped by the phase of the spike in which the channels act. Only significant changes (Benjamini–Hochberg q < 0.05) are numbered. A cell without a number is not significant. An asterisk marks a significant change that may come from ambient RNA. The colour scale is clipped at ±1.5. Values are provided in Extended Data Figure 3-1. Extended Data Figure 3-2 compares the 18-week transcript ratios of BK, Cav3.1, Cav2.1 and IP3R1 with the published staining ratios.

To ask how many Purkinje cells carry the difference, we scored each nucleus along the direction, on the genotype axes, from the average WT mouse to the average KI mouse of its age. The shifted fraction is the fraction of a mouse’s Purkinje nuclei that lie further along this direction than 90 % of the WT nuclei of that age. It is about 0.10 in a WT mouse by construction (Methods). In KI mice, the shifted fraction was 0.535 at 5 weeks and 0.754–0.950 from 18 weeks (Extended Data Figure 2-3), so the difference is carried not by a small subset of cells but by about half of the Purkinje cells at 5 weeks and by most of them from 18 weeks. These values were similar when each mouse was left out in turn and when the direction and the shifted fraction were estimated on separate halves of the counts (count splitting; Neufeld et al. 2024), and the held-out 12-week KI mice, scored without refitting, had shifted fractions of 0.69–1.00 (WT 0.17–0.29; Methods).

RNA from outside the Purkinje nucleus, ambient RNA released during tissue preparation or a second nucleus captured in the same droplet, could also produce this separation. Scores for the marker genes of the other cerebellar cell types did not differ between KI and WT Purkinje nuclei, except for endothelial-cell RNA, which was higher in KI Purkinje nuclei from 18 weeks. The part of each component score carried by the Purkinje cell’s own marker genes separated KI from WT in the same direction in 10 of the 14 component-ages with complete separation, which indicates that the difference is carried by the Purkinje cell’s own transcripts, not only by contamination. It does not show where the change originates, because the knock-in allele is expressed in every cell type.

In KI mice, the nuclei did not converge on one state. Their spread along the components increased with age on 3 of the 4 components (in WT, it decreased on all 4), and at 30 weeks the KI spread exceeded the WT spread on all 4 components, with complete separation on 3. At 24 and 30 weeks, no KI component-age showed two separate groups of nuclei (0 of 8; WT 1 of 8; Methods).

### A loss of potassium and calcium channel transcripts, detected by 12 weeks

Differential expression with the mouse as the unit of analysis returned 2,729 hits at a false discovery rate of 0.05 across 27,415 tests. A hit is one gene in one cell type at one age, and this count covers all cell types. Of these hits, 290 were in Purkinje cells (32 at 5 weeks, 95 at 18 weeks, 70 at 24 weeks and 93 at 30 weeks). To validate our data, we asked whether a previously reported set of genes downregulated in Purkinje cells of SCA1 mice (the Magenta module; Ingram et al. 2016) was also downregulated here. It was significantly downregulated in KI Purkinje cells at 18, 24 and 30 weeks (q = 3 × 10⁻⁴⁰ to 5 × 10⁻³² at the three ages; Extended Data Figure 3-1).

Changes in ion-channel genes fell into four groups (Figure 3; log2 fold change at 18 weeks in parentheses). Calcium channel transcripts were reduced: Cav2.1 (Cacna1a, P/Q-type; −0.57), Cav3.1 (Cacna1g, T-type; −0.62) and the Cav2.1 auxiliary subunit α2δ2 (Cacna2d2; −0.79). Transcripts of potassium channels involved in repolarization of the spike and in shaping the period after it were also reduced: Kv3.3 (Kcnc3; −0.59), BK (Kcnma1; −0.48) and Kv4.3 (Kcnd3; −0.77), as well as the BK auxiliary subunit β4 (Kcnmb4; −1.40, already reduced at 5 weeks) and the Kv4 auxiliary subunits KChIP1 (Kcnip1, from 24 weeks) and DPP10 (Dpp10, at 30 weeks). The auxiliary subunit Kvβ1 (Kcnab1; −0.72), which binds mainly Kv1 channels, was also reduced. The transcripts of two proteins of the calcium store of the endoplasmic reticulum were also reduced: the IP3 receptor IP3R1 (Itpr1; −0.76) and the pump SERCA3 (Atp2a3; −0.92).

The sodium channels Nav1.6 (Scn8a), Nav1.1 (Scn1a) and Nav1.2 (Scn2a) and their regulatory proteins Navβ4 (Scn4b) and FGF14 (Fgf14) showed no significant change between KI and WT at any age, although a change that is not significant can still have a point estimate well away from zero (Navβ4, −0.64 at 18 weeks). Among the sodium-channel transcripts, only that of the auxiliary subunit Navβ1 (Scn1b) was reduced, at 18 weeks. The low-threshold potassium channels Kv1.1 (Kcna1) and Kv1.2 (Kcna2) also showed no significant change. The pacemaker channel HCN1 (Hcn1) increased at 5 weeks only, and the M-current channels Kv7.2, Kv7.3 and Kv7.5 (Kcnq2, Kcnq3 and Kcnq5) and the small-conductance calcium-activated potassium channel SK2 (Kcnn2) showed no significant change. In the discovery mice, the transcripts of the five pore-forming subunits (Kv3.3, BK, Kv4.3, Cav2.1 and Cav3.1) were reduced at 18, 24 and 30 weeks (by 28–41 %, 17–36 % and 24–36 %), but not at 5 weeks. These are group-level changes, estimated from per-mouse pseudobulk. Whether the five reductions occur together within individual nuclei is a separate question that we did not test.

The same reduction was found in the six held-out 12-week mice, which were not used to find these genes. In KI Purkinje cells at 12 weeks, the transcripts of the five channels were reduced (q < 0.05 for each): Kv3.3 (log2 fold change −0.55), Cav3.1 (−0.43), Cav2.1 (−0.40), Kv4.3 (−0.39) and BK (−0.27), as was SERCA3 (−0.51, q = 0.007). IP3R1 was reduced by a similar amount but did not reach significance (−0.32, q = 0.10; Figure 4). Thus, the reduction of the channel transcripts was first detected at 12 weeks.

**Figure 4.**
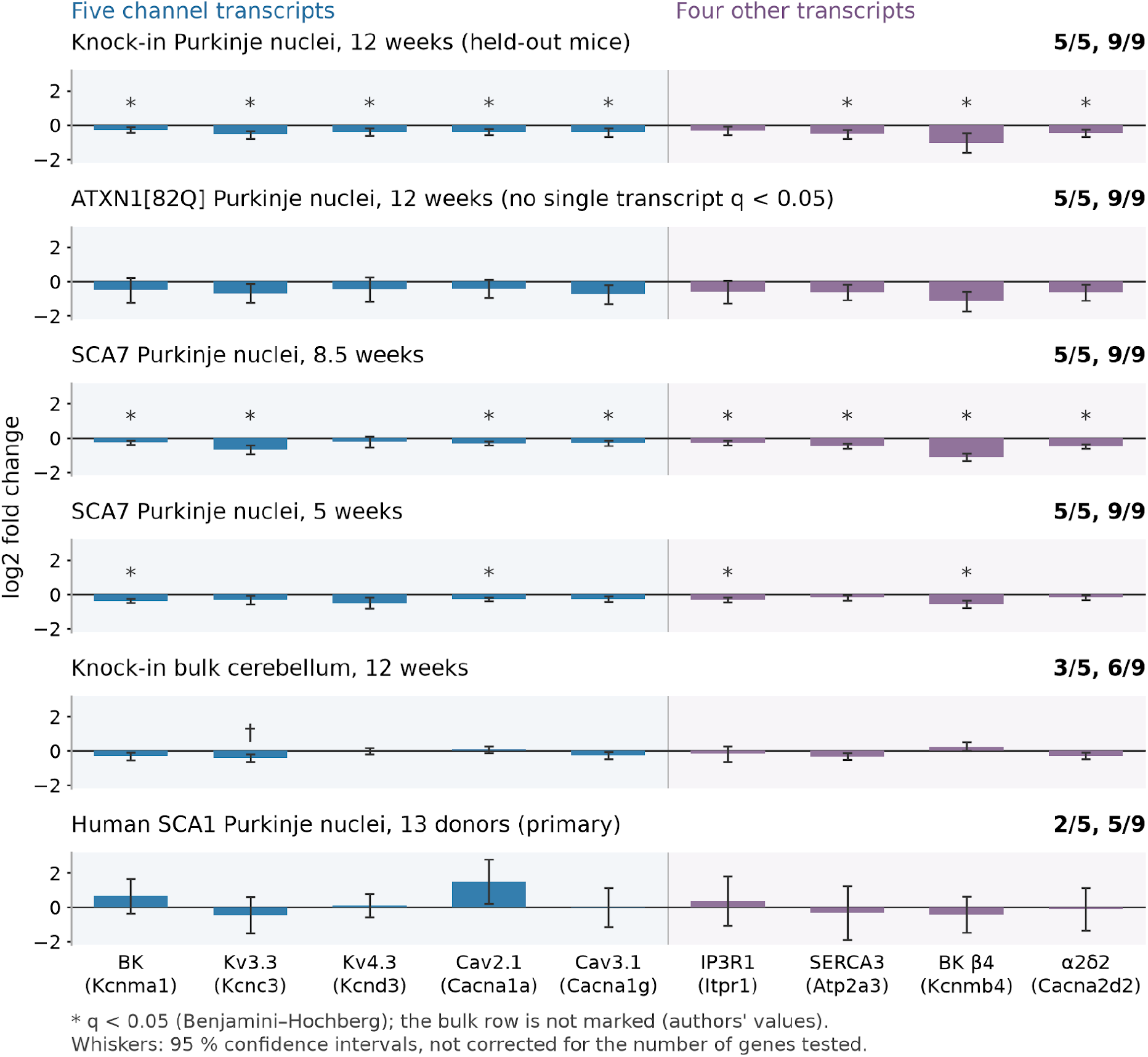
Replication of the channel transcript reduction in other datasets. Log2 fold change (KI − WT, or disease − control) of the five channel transcripts (blue) and the four other transcripts of the set (purple) in six comparisons, from top to bottom: 12-week mice held out of the discovery analyses; ATXN1[82Q] Purkinje nuclei at 12 weeks; SCA7 Purkinje nuclei at 8.5 weeks and at 5 weeks; bulk knock-in cerebellum at 12 weeks; and human SCA1 Purkinje nuclei (13 donors). Transcripts are labelled with the channel or protein and, in parentheses, its gene. Bars are estimates and whiskers are 95 % confidence intervals (for the bulk data the whiskers are approximate, derived from the authors’ p values). The number of negative estimates among the five channel transcripts and among all nine transcripts is shown to the right of each row. These counts give the direction of the estimates, not their significance. All five channel transcripts reached q < 0.05 in the held-out mice (Extended Data Figure 3-1), and no single transcript did in the ATXN1[82Q] data. Asterisks mark the transcripts with q < 0.05 (Benjamini–Hochberg). In the held-out mice, q is taken over the 499 tests of Purkinje nuclei at 12 weeks in the five-age run (Extended Data Figure 3-1), and in the other single-nucleus datasets over all genes tested in that dataset (9,760 for ATXN1[82Q], 12,111 for SCA7 and 7,008 for human SCA1, Extended Data Figure 4-1). The bulk row is not marked because its values are the authors’. Whiskers are not corrected for the number of genes tested, whereas q values are. The SCA7 rows are exploratory because they concern another disease gene and another mouse model. † In the bulk data, the Kv3.3 (Kcnc3) value is the authors’ value for the merged Kcnc3,Napsa locus. Values are provided in Extended Data Figure 4-1.

We then asked whether the reduction is reproduced in outside datasets (Figure 4; Extended Data Figure 4-1), counting in each how many of the five channel transcripts, and of nine transcripts (the five channels plus IP3R1, SERCA3, BK β4 and α2δ2), were lower in disease than in controls. In single-nucleus RNA-seq of Pcp2-ATXN1[82Q] mice, which express expanded human ATXN1 in Purkinje cells only (Borgenheimer et al. 2022; three mice per genotype, 12 weeks), all five channel transcripts and all nine transcripts were lower than in WT (median log2 fold change of the five channel transcripts −0.51; BK β4 −1.16), although with three mice per group no single transcript reached q < 0.05. In an exploratory analysis of SCA7 266Q mice, which carry the expansion in Atxn7 rather than Atxn1 (four mice per genotype), all nine transcripts were also lower in Purkinje-enriched nuclei at 5 and 8.5 weeks.

In human SCA1 cerebellum (the human arm of GSE246183), only 13 of the 20 donors (7 control, 6 SCA1) had at least 10 Purkinje nuclei. In these donors, two of the five channel transcripts, and five of the nine transcripts in total, were lower in SCA1, and Cav2.1 (CACNA1A) was higher (+1.51, nominal p = 0.025). With so few nuclei per donor, we consider this result inconclusive rather than negative. In bulk data from 12-week knock-in cerebellum (Driessen et al. 2018), 3 of the 5 channel transcripts were decreased, but a control set of 113 Purkinje-enriched genes was reduced as much (median −0.30 vs −0.27), and in ATXN1[82Q] cerebellum (Ingram et al. 2016) the control set was reduced even more than the channel transcripts (Methods). Because the nine transcripts change together and are not independent, the sign-test p values are nominal (Methods), and we report the counts of direction instead.

### SCA1 knock-in Purkinje cells have more variable interspike intervals

To test whether the transcript changes suffice, we used the published Purkinje cell model of Zang et al. (2018) and its extension with background synaptic input (Zang and De Schutter 2021), in which the conductances can be scaled by the KI/WT transcript ratios and then restored singly or in combination. The firing rate, the variability of the interspike interval and the restoration of Cav2.1 were also tested in a second published model (Masoli et al. 2015; Methods). The target was the firing of Purkinje cells of the same knock-in line recorded at 14 weeks in acute slices of the anterior cerebellum and the nodular zone (Bushart et al. 2021; values in Table 2 and Methods). Their firing rate did not differ significantly from that of WT in either region: the KI/WT ratio of the group-mean firing rates, where 1 means no difference, was 0.86 in the nodular zone and 1.33 in the anterior cerebellum, and we use the span between these two values as the recorded rate range. Their interspike intervals were more variable: the KI/WT ratio of the group-mean CVs (the CV ratio) was 1.8 in the anterior cerebellum and 2.9 in the nodular zone. The higher CV persisted in the presence of synaptic blockers, which suggests that it is intrinsic to the Purkinje cell. Each run with the conductances scaled by the KI/WT ratios (a KI-scaled run) was compared with its own WT run on the four readouts of Table 2: the firing rate ratio, the CV ratio, the afterhyperpolarization difference and the block-threshold ratio.

**Table 2.**
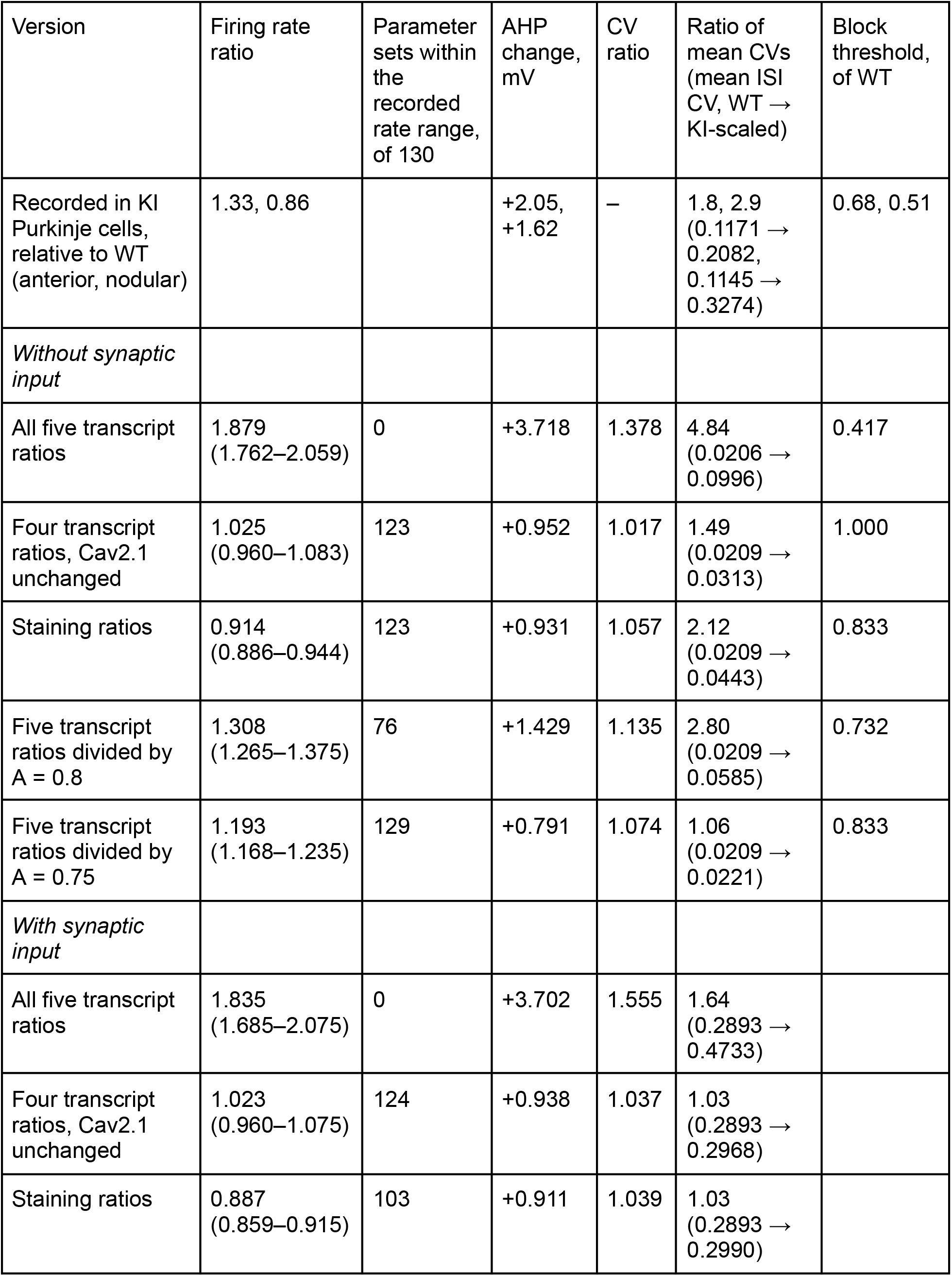
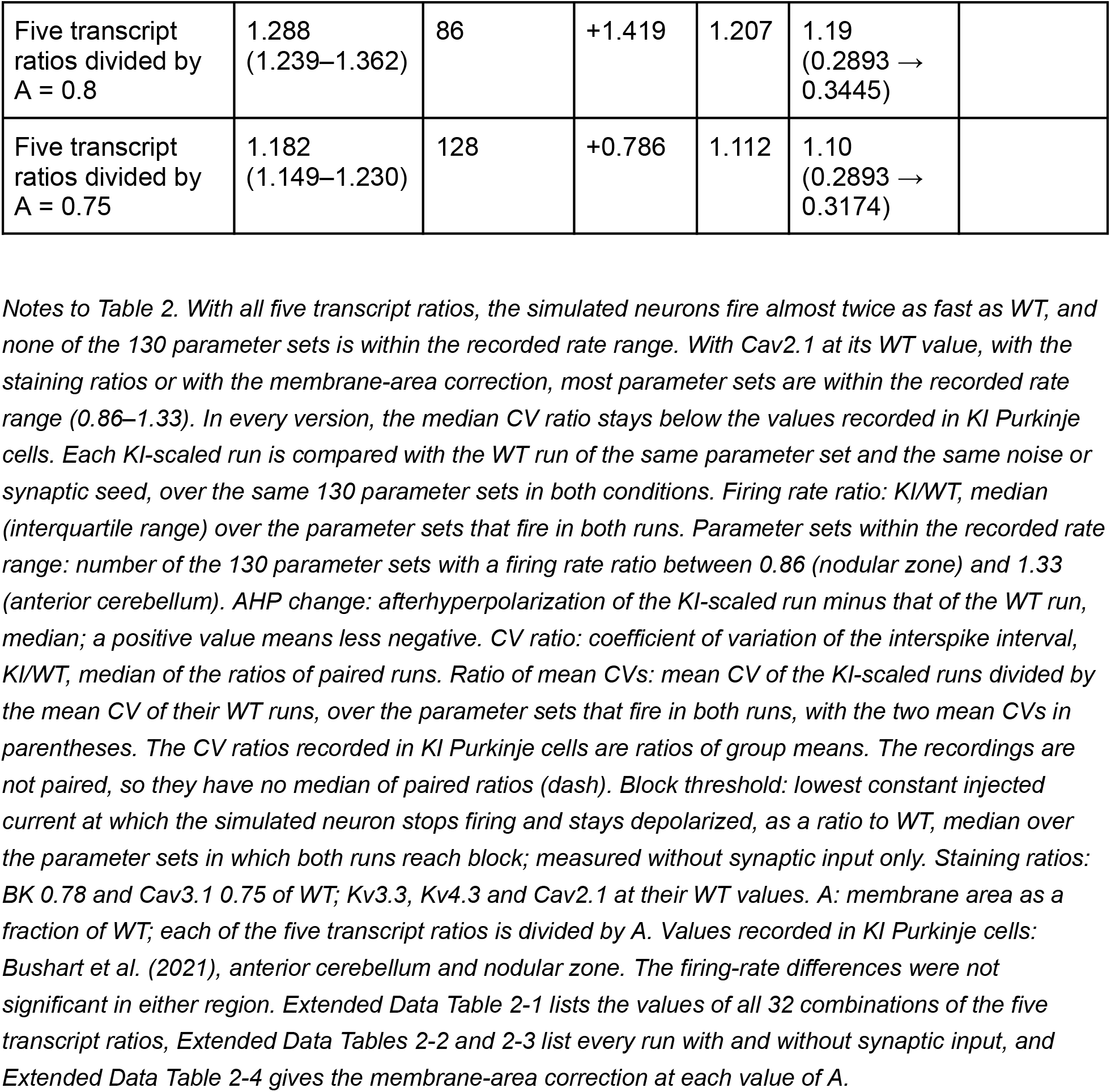
Simulated neurons compared with the phenotype recorded in KI Purkinje cells, without and with synaptic input.

| Version | Firing rate ratio | Parameter sets within the recorded rate range, of 130 | AHP change, mV | CV ratio | Ratio of mean CVs (mean ISI CV, WT → KI-scaled) | Block threshold, of WT |
|---|---|---|---|---|---|---|
| Recorded in KI Purkinje cells, relative to WT (anterior, nodular) | 1.33, 0.86 |  | +2.05, +1.62 | – | 1.8, 2.9 (0.1171 → 0.2082, 0.1145 → 0.3274) | 0.68, 0.51 |
| *Without synaptic input* |  |  |  |  |  |  |
| All five transcript ratios | 1.879 (1.762–2.059) | 0 | +3.718 | 1.378 | 4.84 (0.0206 → 0.0996) | 0.417 |
| Four transcript ratios, Cav2.1 unchanged | 1.025 (0.960–1.083) | 123 | +0.952 | 1.017 | 1.49 (0.0209 → 0.0313) | 1.000 |
| Staining ratios | 0.914 (0.886–0.944) | 123 | +0.931 | 1.057 | 2.12 (0.0209 → 0.0443) | 0.833 |
| Five transcript ratios divided by A = 0.8 | 1.308 (1.265–1.375) | 76 | +1.429 | 1.135 | 2.80 (0.0209 → 0.0585) | 0.732 |
| Five transcript ratios divided by A = 0.75 | 1.193 (1.168–1.235) | 129 | +0.791 | 1.074 | 1.06 (0.0209 → 0.0221) | 0.833 |
| *With synaptic input* |  |  |  |  |  |  |
| All five transcript ratios | 1.835 (1.685–2.075) | 0 | +3.702 | 1.555 | 1.64 (0.2893 → 0.4733) |  |
| Four transcript ratios, Cav2.1 unchanged | 1.023 (0.960–1.075) | 124 | +0.938 | 1.037 | 1.03 (0.2893 → 0.2968) |  |
| Staining ratios | 0.887 (0.859–0.915) | 103 | +0.911 | 1.039 | 1.03 (0.2893 → 0.2990) |  |
| Five transcript ratios divided by A = 0.8 | 1.288 (1.239–1.362) | 86 | +1.419 | 1.207 | 1.19 (0.2893 → 0.3445) |  |
| Five transcript ratios divided by A = 0.75 | 1.182 (1.149–1.230) | 128 | +0.786 | 1.112 | 1.10 (0.2893 → 0.3174) |  |

*Notes to Table 2. With all five transcript ratios, the simulated neurons fire almost twice as fast as WT, and none of the 130 parameter sets is within the recorded rate range. With Cav2.1 at its WT value, with the staining ratios or with the membrane-area correction, most parameter sets are within the recorded rate range (0.86–1.33). In every version, the median CV ratio stays below the values recorded in KI Purkinje cells. Each KI-scaled run is compared with the WT run of the same parameter set and the same noise or synaptic seed, over the same 130 parameter sets in both conditions. Firing rate ratio: KI/WT, median (interquartile range) over the parameter sets that fire in both runs. Parameter sets within the recorded rate range: number of the 130 parameter sets with a firing rate ratio between 0.86 (nodular zone) and 1.33 (anterior cerebellum). AHP change: afterhyperpolarization of the KI-scaled run minus that of the WT run, median; a positive value means less negative. CV ratio: coefficient of variation of the interspike interval, KI/WT, median of the ratios of paired runs. Ratio of mean CVs: mean CV of the KI-scaled runs divided by the mean CV of their WT runs, over the parameter sets that fire in both runs, with the two mean CVs in parentheses. The CV ratios recorded in KI Purkinje cells are ratios of group means. The recordings are not paired, so they have no median of paired ratios (dash). Block threshold: lowest constant injected current at which the simulated neuron stops firing and stays depolarized, as a ratio to WT, median over the parameter sets in which both runs reach block; measured without synaptic input only. Staining ratios: BK 0.78 and Cav3.1 0.75 of WT; Kv3.3, Kv4.3 and Cav2.1 at their WT values. A: membrane area as a fraction of WT; each of the five transcript ratios is divided by A. Values recorded in KI Purkinje cells: Bushart et al. (2021), anterior cerebellum and nodular zone. The firing-rate differences were not significant in either region. Extended Data Table 2-1 lists the values of all 32 combinations of the five transcript ratios, Extended Data Tables 2-2 and 2-3 list every run with and without synaptic input, and Extended Data Table 2-4 gives the membrane-area correction at each value of A.*

### In the Zang models, scaling all five conductances by their transcript ratios increases both the median firing rate and the CV

To test whether the change in firing of KI Purkinje cells can be explained by the change in transcripts, we multiplied the maximum conductance of each of the five channels by its KI/WT transcript ratio at 18 weeks (Kv3.3 0.667, BK 0.719, Kv4.3 0.588, Cav2.1 0.672, Cav3.1 0.649). For the two calcium channels, the scaled parameter of the model mechanism is the maximum permeability, which we include under “conductance” throughout. Table 3 summarizes what each of these channels does in Purkinje cells. Of the four ages of the discovery mice, we chose 18 weeks because it is closest to the age at which the KI Purkinje cells were recorded (14 weeks). Applying the 24-week or 30-week ratios to the published parameter set (five noise seeds) also raised the firing rate, to 1.53 and 1.51 times the WT rate, against 1.91 with the 18-week ratios (Extended Data Figure 5-1). We applied the 18-week ratios in the Purkinje cell model of Zang et al. (2018) and in its successor with background synaptic input (Zang and De Schutter 2021). Both models have a full dendritic morphology and are simulated at 34 °C. In such a model, different combinations of conductances can lead to the same firing, so a result obtained with one parameter set may not hold for another. We therefore drew 200 parameter sets by scaling eight groups of conductances by factors between 0.67 and 1.5 of their published values, and retained the 130 parameter sets that fired tonically with the WT conductances and without synaptic input. We ran all 130 parameter sets without synaptic input (the 2018 model with a somatic noise current) and with background synaptic input (the 2021 model). Each KI-scaled run was compared with the WT run of the same parameter set and the same noise or synaptic seed.

**Table 3.** What each of the five channels does in Purkinje cells, and how it changes in KI mice and in the model.

| Channel (mouse gene; human gene) | Type | Role in Purkinje cells | KI/WT transcript ratio, 18 weeks | KI/WT staining ratio (anterior, nodular) | Reduced alone in the model: firing rate ratio, CV ratio |
|---|---|---|---|---|---|
| Kv3.3 (Kcnc3; KCNC3) | Voltage-gated potassium channel of the Kv3 family; opens only at strongly depolarized potentials, and opens and closes fast | Potassium currents blocked by low concentrations of TEA, as Kv3 currents are, repolarize each spike quickly and close soon after it, which supports high firing rates (Raman and Bean 1999). At the soma, Kv3.3 is needed for the repeated spikelets of the complex spike that a climbing fibre evokes (Zagha et al. 2008). | 0.667 | Not measured | 0.93, 0.94 |
| BK (Kcnma1; KCNMA1) | Potassium channel of large conductance, opened by calcium and by depolarization; also called KCa1.1 or Slo1 | Opened by calcium entering through Cav2.1 channels; shapes the spike and the afterhyperpolarization that follows it (Edgerton and Reinhart 2003; Womack et al. 2004). Mice lacking BK are ataxic, and their Purkinje cells fire much less, because depolarization inactivates spike generation (Sausbier et al. 2004). | 0.719 | 0.79, 0.77 | 1.07, 1.05 |
| Kv4.3 (Kcnd3; KCND3) | Voltage-gated A-type potassium channel; activates at low voltages and inactivates fast | Present in the dendritic shafts and spines. Its current limits the calcium spikes that a climbing fibre evokes in the dendrites; when these channels are inactivated, the calcium spikes spread into the distal dendrites (Otsu et al. 2014). | 0.588 | Not measured | 1.28, 0.98 |
| Cav2.1 (Cacna1a; CACNA1A) | Calcium channel activated at high voltages; P/Q-type (P-type in Purkinje cells) | Present in the dendrites and spines (Kulik et al. 2004; Indriati et al. 2013). It carries the dendritic calcium spikes (Otsu et al. 2014), and the calcium entering through it opens BK and SK potassium channels (Womack et al. 2004). In mice with a Cav2.1 mutation, the pacemaking of Purkinje cells is less precise, which causes ataxia (Walter et al. 2006). | 0.672 | 1.01, 0.93 | 1.75, 1.40 |
| Cav3.1 (Cacna1g; CACNA1G) | Calcium channel activated at low voltages;<br>T-type | Concentrated in the dendritic spines at parallel fibre synapses, where it is coupled to the metabotropic glutamate receptor mGluR1 and enhanced by parallel fibre input (Isope et al. 2012). Under resting conditions, T-type channels carry the small calcium signals that a climbing fibre evokes in the dendrites (Otsu et al. 2014). | 0.649 | 0.77, 0.73 | 0.82, 1.02 |

*Notes to Table 3. Role: as reported in Purkinje cells of rats and mice; not measured in this study. KI/WT transcript ratio: in Purkinje nuclei at 18 weeks, the factor by which each conductance was scaled in the model. KI/WT staining ratio: staining intensity in the molecular layer of the anterior cerebellum and the nodular zone (Bushart et al. 2021). BK and Cav3.1 staining was significantly reduced, and Cav2.1 staining was not. Reduced alone in the model: this conductance scaled by its transcript ratio and the other four left unchanged, in the Zang et al. (2018) model without synaptic input; firing rate ratio and CV ratio, KI/WT, median over the parameter sets that fire in both runs (Extended Data Table 2-1). SK: small-conductance calcium-activated potassium channel. TEA: tetraethylammonium.*

Without synaptic input, scaling all five conductances raised the firing rate (Figure 5A, B; Table 2). Two of the 130 parameter sets went into depolarization block and the remaining 128 still fired, with a median firing rate ratio of 1.879 (interquartile range 1.762–2.059). In none of the 130 parameter sets was the firing rate ratio within the range recorded in Purkinje cells of Atxn1 154Q/2Q mice, from 0.86 in the nodular zone to 1.33 in the anterior cerebellum (Bushart et al. 2021). The CV ratio was 1.378 (KI Purkinje cells: 1.8 anterior, 2.9 nodular), and the afterhyperpolarization was 3.718 mV less negative than in WT (KI Purkinje cells: 2.05 mV anterior, 1.62 mV nodular).

**Figure 5.**
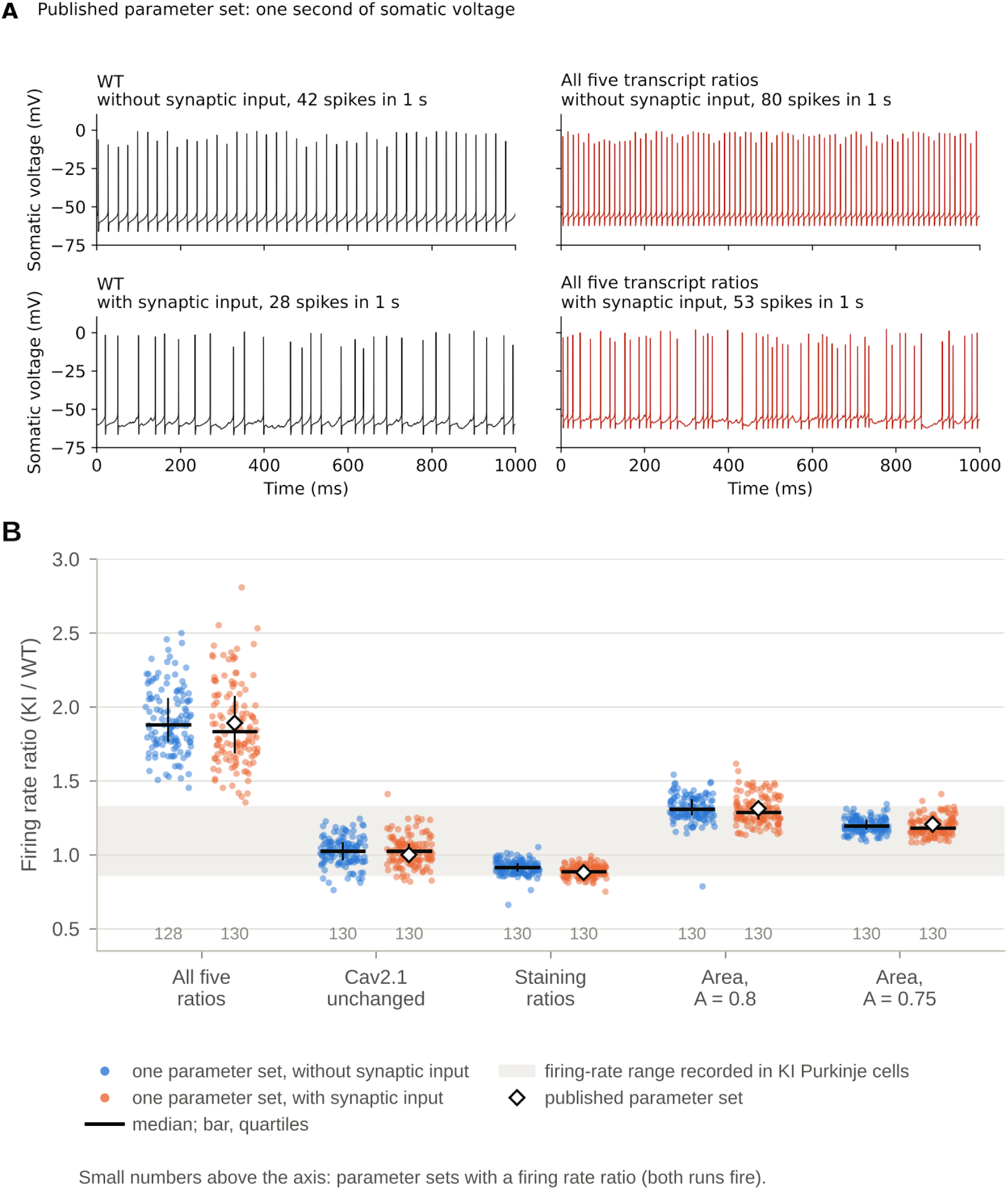
Simulated neurons scaled by the KI/WT transcript ratios, compared with the firing of KI Purkinje cells. (A) Somatic membrane voltage of the published parameter set over 1 second (as in a whole-cell current-clamp recording; the firing rate and CV of KI Purkinje cells were recorded cell-attached, from spike times only), for WT and with all five transcript ratios, without synaptic input (42 and 80 spikes) and with synaptic input (28 and 53 spikes). (B) Firing rate ratio (KI/WT) of each of the 130 parameter sets in five versions. All five transcript ratios: Kv3.3 0.667, BK 0.719, Kv4.3 0.588, Cav2.1 0.672 and Cav3.1 0.649 of WT. Cav2.1 unchanged: Kv3.3, BK, Kv4.3 and Cav3.1 at their transcript ratios; Cav2.1 unchanged. Staining ratios: BK 0.78 and Cav3.1 0.75 of WT; Kv3.3, Kv4.3 and Cav2.1 unchanged. Membrane-area correction: each of the five transcript ratios divided by A, the membrane area as a fraction of WT, with A = 0.8 and A = 0.75. Blue, without synaptic input; orange, with synaptic input; line, median; bar, quartiles; diamond, published parameter set; shaded band, the recorded rate range (0.86–1.33); small numbers, number of parameter sets with a ratio. Extended Data Figure 5-1 gives every run of the published parameter set (five noise seeds) with the transcript ratios of 18, 24 and 30 weeks, with the membrane-area correction and with the α2δ2 and BK β4 changes. Extended Data Figure 5-2 shows one second of somatic voltage of the published parameter set with synaptic input for WT and the five versions of panel B.

With synaptic input, the result was the same (Figure 5A, B; Table 2). All 130 parameter sets kept firing, and none went into depolarization block. The firing rate ratio was 1.835 (interquartile range 1.685–2.075), and again none of the 130 parameter sets was within the recorded rate range. The CV ratio was 1.555, and the afterhyperpolarization was less negative by 3.702 mV. The published parameter set scaled with all five transcript ratios (diamonds in Figure 5B), run separately with synaptic input, gave a firing rate ratio of 1.891, a CV ratio of 1.609 and an afterhyperpolarization less negative by 3.864 mV. Scaling all five conductances by their transcript ratios therefore increased both the median firing rate and the CV of the simulated neurons. Extended Data Figure 5-2 shows one second of somatic voltage of the published parameter set with synaptic input for WT and the five versions of Figure 5B.

### The Cav2.1 reduction accounts for most of the firing-rate increase in the model

One of the five transcript ratios does not match the staining in KI mice: the Cav2.1 transcript was 0.672 of WT, but Cav2.1 staining of the molecular layer was not significantly different from WT (Bushart et al. 2021; Extended Data Figure 3-2). To find which reductions are responsible for the faster firing, we ran all 32 combinations in which each of the five conductances was either scaled by its 18-week transcript ratio or left at its WT value. We call each combination a version, and all versions used the same 130 parameter sets and seeds (Methods; Extended Data Table 2-1). Reduced alone, Cav2.1 raised the firing rate the most (median firing rate ratio 1.75), Kv4.3 and BK less (1.28 and 1.07), and Kv3.3 and Cav3.1 lowered it (0.93 and 0.82). Only the Cav2.1 reduction raised the CV ratio markedly (1.40, against 0.94–1.05 for the other four). The effect of each reduction hardly depended on which of the others were reduced, and with several reductions together, the firing rate ratio was close to the product of the ratios of each reduction alone (Methods).

With all five transcript ratios, the simulated neurons fired at a median of 1.88 times the WT rate, with none of the 130 parameter sets within the recorded rate range, and the Cav2.1 reduction accounted for most of this increase: in each of the 16 combinations of the other four conductances, reducing Cav2.1 raised the median firing rate 1.7- to 1.8-fold (Figure 6A). With Cav2.1 left at its WT value and the other four at their transcript ratios, the median firing rate ratio was 1.03 (quartiles 0.96–1.08), with 123 of the 130 parameter sets within the recorded rate range. With synaptic input, it was 1.02, with 124 of the 130 (Figure 5B; Table 2). In this version, without and with synaptic input, the CV ratio stayed close to 1 (1.02 and 1.04), and the afterhyperpolarization was 0.95 and 0.94 mV less negative than in WT, about half of the change recorded in KI Purkinje cells (2.05 mV in the anterior cerebellum and 1.62 mV in the nodular zone). In a second published Purkinje cell model (Masoli et al. 2015), the change went in the same direction but was smaller: the five transcript ratios raised the median firing rate only 1.14-fold at 18 weeks, which is within the recorded rate range, and none of its 14 parameter sets had a higher ISI CV than with its WT settings, but restoring Cav2.1 alone lowered the firing rate in 14 of the 14 (Methods; Extended Data Figure 6-1). In both models, the Cav2.1 reduction accounts for most of the faster firing, and neither reproduces the higher CV of KI Purkinje cells. The size of the rate increase depends on the model: the 1.9-fold increase belongs to the Zang models.

**Figure 6.**
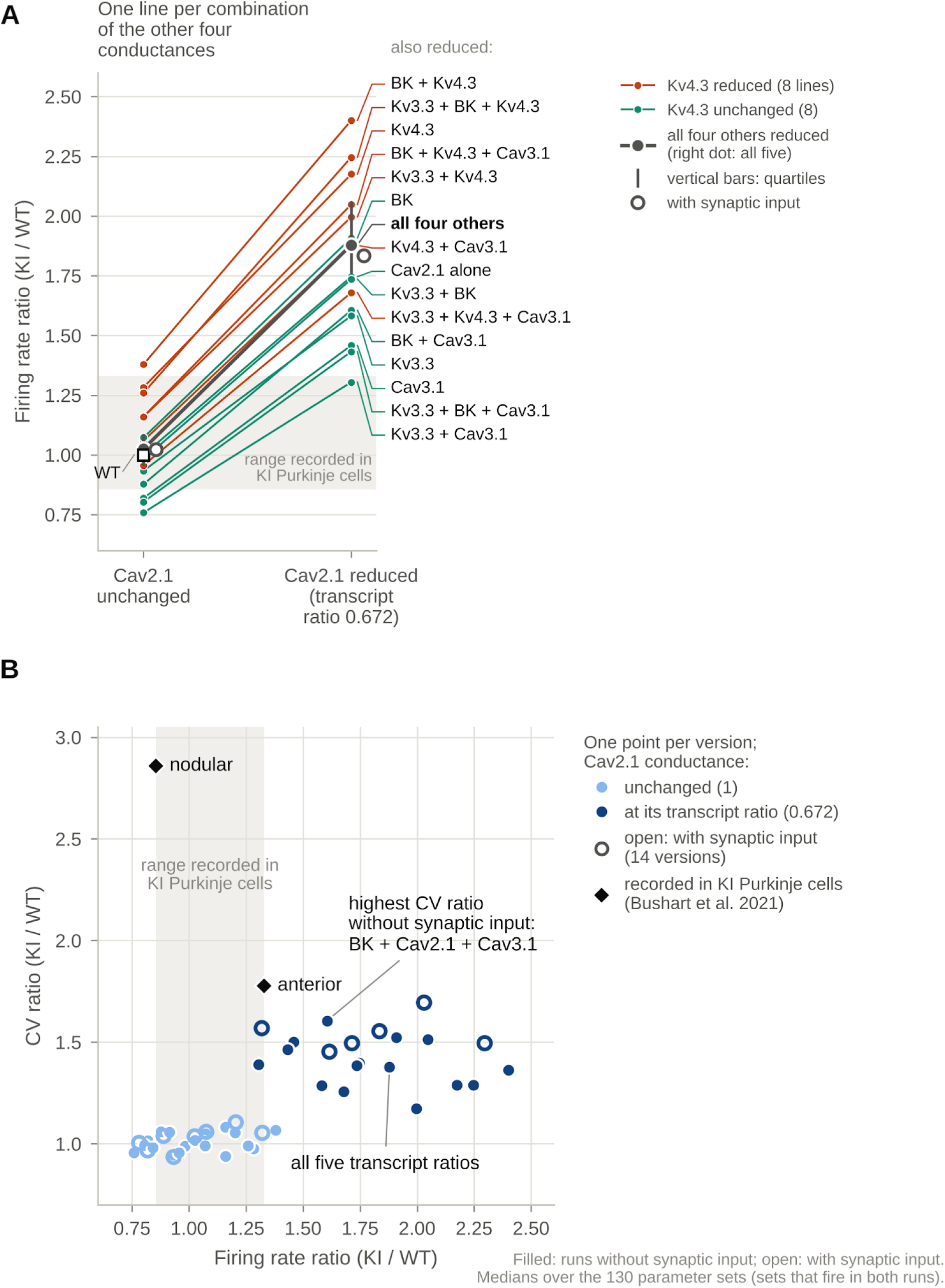
The Cav2.1 reduction makes the simulated neurons fire faster, and in every version the median CV ratio stays below that of KI Purkinje cells. (A) Firing rate of the simulated neurons relative to WT (firing rate ratio, KI/WT) with Cav2.1 unchanged (left dots) and reduced to its transcript ratio (right dots). Each line is one of the 16 combinations of the other four conductances, and the name at the right end of the line indicates which of them are also reduced. Red, Kv4.3 is among them; green, it is not. Each name is joined to its line by a connector of the same colour. Thick grey line, all four others reduced, so that its right dot is the version with all five transcript ratios; vertical bars on this line, quartiles; open circles, the same two versions with synaptic input; square, WT. Quartiles of all 32 versions are given in Extended Data Table 2-1. Reducing Cav2.1 increases the firing rate of the simulated neurons, whatever else is reduced. (B) CV ratio against firing rate ratio. Each point is one version (one setting of the five conductances) and shows the median over the 130 parameter sets. Its colour shows whether the Cav2.1 conductance is unchanged or at its transcript ratio. Filled points, 35 versions without synaptic input; open points, 14 versions with synaptic input (listed in Methods). Diamonds, KI/WT ratios of the group means recorded in KI Purkinje cells (Bushart et al. 2021). Within the recorded rate range, the highest median CV ratio is 1.39 without and 1.57 with synaptic input, against 1.8 and 2.9 recorded in KI Purkinje cells. Shaded band in both panels, the recorded rate range (0.86–1.33). Values are medians over the parameter sets that fire in both runs. Extended Data Figure 6-2 gives the runs with the Cav2.1 conductance lowered in steps on top of the staining ratios, Extended Data Figure 6-3 gives the Kv3.3 and Kv4.3 reductions added to the staining ratios, and Extended Data Figures 6-1 and 6-4 give the runs in the model of Masoli et al. (2015).

### Staining ratios move the afterhyperpolarization and the block threshold towards the recordings, but by less

We then replaced the transcript ratios with the staining ratios, the KI/WT staining intensities of the molecular layer averaged over the two regions (BK 0.78, Cav3.1 0.75; Bushart et al. 2021), and left the other three conductances unchanged (Table 2). The simulated neurons fired slightly slower than WT (median firing rate ratio 0.91; 123 of the 130 parameter sets within the recorded rate range), and their afterhyperpolarization and block threshold moved in the direction recorded in KI Purkinje cells, but by less (afterhyperpolarization 0.93 mV less negative; block threshold 0.83 of WT). Staining of the molecular layer measures Cav2.1 protein in the dendrites and spines of Purkinje cells together with that in the parallel fibre terminals of granule cells (Kulik et al. 2004; Indriati et al. 2013). It also counts channel protein inside the cell as well as that in the membrane, so it does not show how many of the channels are in the Purkinje cell membrane. It therefore cannot exclude a partial loss of Cav2.1 from that membrane.

### In every version, the median CV ratio stays below that of KI Purkinje cells

In every version, the median CV ratio stayed below that of KI Purkinje cells, whose CV was 1.8 and 2.9 times that of WT (Figure 6B; Extended Data Tables 2-2 and 2-3). Among the versions whose median firing rate lay within the recorded rate range, the highest median CV ratio was 1.39 without synaptic input (Kv3.3, Cav2.1 and Cav3.1 at their transcript ratios) and 1.57 with synaptic input (Kv4.3 unchanged and the other four at their transcript ratios). Higher median CV ratios, up to 1.61 without and 1.70 with synaptic input, came only with firing well above the WT rate (firing rate ratios 1.61 and 2.03). These statements concern the median of each version across the 130 parameter sets, not every simulated neuron. With synaptic input, 23 runs in 7 of the 21 versions had a firing rate ratio within the recorded rate range and a CV ratio between 1.8 and 2.9, 21 of them with pauses, and without synaptic input 2 runs did so (Extended Data Tables 2-2 and 2-3). Falling within both ranges does not make a run match either region, or the four readouts together.

These CV ratios are medians of the ratios of paired runs, whereas the values recorded in KI Purkinje cells are ratios of group means, and the two summaries differ when a few simulated neurons pause (Table 2). Without synaptic input, the ratio of the mean CVs reached 1.8 or more in 13 of the 24 versions whose median firing rate lay within the recorded rate range, each time because 1 to 10 of the 130 parameter sets paused. Even then, the mean CV of the KI-scaled runs (0.04–0.11) stayed below the mean CVs recorded in KI Purkinje cells (0.21 and 0.33), and that of the WT runs was 0.02 (Extended Data Table 2-3). With synaptic input, the mean CV of the WT runs was 0.29, and the ratio of the mean CVs was at most 1.54 in the versions whose median firing rate lay within the recorded rate range (Extended Data Table 2-2). The 130 parameter sets are not a sample of real cells, so the model shows that rare pauses can produce a ratio of means as large as that recorded, not how often this happens in KI Purkinje cells.

### A correction for the smaller membrane brings the median firing rate into the recorded rate range but does not reproduce the higher CV

Purkinje cells of KI mice also have a smaller membrane than those of WT mice. Their capacitance was 0.77 of WT at 6–11 weeks (whole cell; 0.75 for the dendritic tree; Watase et al. 2002) and 0.80 of WT at 12 weeks (Shuvaev et al. 2022). On a smaller membrane, the same number of channels gives a higher conductance density, so we divided each of the five transcript ratios by A, the membrane area as a fraction of WT, changing only the conductance densities and not the shape of the simulated neuron (Extended Data Table 2-4). At A = 0.80, the median firing rate ratio fell from 1.88 to 1.31, and 76 of the 130 parameter sets were within the recorded rate range (0.86–1.33). At A = 0.75, it fell to 1.19, still above the WT rate, with 129 of the 130 parameter sets within that range. The CV ratio fell with the firing rate, to 1.135 at A = 0.80 and 1.074 at A = 0.75, and at A = 0.75 the afterhyperpolarization change was less than half of that recorded in KI Purkinje cells (Table 2). The membrane-area correction can therefore bring the median firing rate of the simulated neurons into the recorded rate range, but it does not reproduce the higher CV.

## Discussion

We analysed public single-nucleus RNA-seq of Atxn1 154Q/2Q knock-in (KI) and WT cerebellum. KI Purkinje nuclei were reproducibly shifted from 5 weeks, and the transcripts of five ion channels, Kv3.3, BK, Kv4.3, Cav2.1 and Cav3.1, were reduced from 18 weeks in the discovery mice and at 12 weeks in the held-out mice, and were lower, though not significantly for any single transcript, in ATXN1[82Q] Purkinje nuclei. Applied as conductance ratios in a published Purkinje cell model, these KI/WT transcript ratios made the simulated neurons fire at a median of 1.88 times the WT rate. The firing rate came into the recorded rate range when Cav2.1 was kept at its WT value, with the staining ratios or with a correction for the smaller membrane. The variability of the interspike interval did not: among all versions with a median firing rate in that range, the median CV ratio was at most 1.39 without and 1.57 with synaptic input, against 1.8 and 2.9 in KI Purkinje cells (Figure 6B). The ratio of the mean CVs, the summary used for the recordings, reached 1.8 or more in some of these versions without synaptic input, but only because a few simulated neurons paused, and even then the mean CV of the KI-scaled runs stayed below that of KI Purkinje cells (Table 2; Extended Data Table 2-3). Thus, scaling the channel conductances by their transcript changes could bring the firing rate of the simulated neurons into the recorded rate range, but the median increase in the variability of the interspike interval remained smaller than that recorded in KI Purkinje cells.

The transcript ratios cannot be read as conductance ratios. Between a transcript and a conductance lie translation, assembly with auxiliary subunits, transport to the membrane and gating, and the size of a change in mRNA may not carry over to protein (Schwanhäusser et al. 2011; Liu et al. 2016). In cortical cells that were recorded and then sequenced (Gouwens et al. 2020; Scala et al. 2021), channel-gene expression predicted the fitted parameters of single-compartment models only in part (cross-validated R² = 0.17), some conductances better than others (Bernaerts et al. 2025). In KI mice, the Cav2.1 transcript is reduced by a third, but Cav2.1 staining of the molecular layer is not significantly reduced, whereas BK and Cav3.1 staining fell with their transcripts (Bushart et al. 2021; Extended Data Figure 3-2). Staining, in turn, does not show how many channels are in the membrane or how much current they carry. In heterozygous Cacna1a D1585N knock-in mice, Cav2.1 staining was similar to that of WT, whereas the peak Cav2.1 current density of Purkinje cells dissociated from 11- to 20-day-old mice was about 30 % lower (Niu et al. 2026). Our simulations therefore test the transcript changes, not the channels of KI Purkinje cells.

Cav2.1 shows the problem most clearly. In the model, reducing Cav2.1 raised the median firing rate whatever else was reduced (Figure 6A). In mouse Purkinje cells, a partial loss of Cav2.1 (P/Q-type) current can make the interspike intervals more variable: in tottering mice, which carry a mutation in Cav2.1 that lowers the P/Q-type current density of their Purkinje cells to about 60 % of WT (Wakamori et al. 1998), the CV was 2.7 times that of WT at a similar firing rate (Alviña and Khodakhah 2010). In SCA6 84Q/84Q mice, the CV was 1.6 times that of WT, at a lower firing rate (Jayabal et al. 2016), whereas in rat slices, blocking about half of the P/Q-type channels raised both the firing rate and the CV (Womack et al. 2004). Either the loss of the Cav2.1 transcript has not reached the channel or it has, and the model lacks what makes mouse Purkinje cells respond to it with more variable intervals rather than faster firing. The staining cannot tell these apart, because the molecular layer also contains the Cav2.1 of parallel fibre terminals (Kulik et al. 2004; Indriati et al. 2013), and to our knowledge the Cav2.1 current of KI Purkinje cells has not been measured.

A smaller membrane is the second way to recover the firing rate. Dell’Orco et al. (2015) suggested that, with fewer channels, a smaller membrane keeps channel density closer to normal. In 154Q/2Q mice, Purkinje cell capacitance was 0.77 of WT at 6–11 weeks (Watase et al. 2002) and 0.80 of WT at 12 weeks (Shuvaev et al. 2022). With the five transcript ratios divided by a membrane area of 0.80 of WT, without synaptic input, the median firing rate ratio fell from 1.88 to 1.31, with 76 of the 130 parameter sets in the recorded rate range, but the median CV ratio fell too, to 1.135 (Extended Data Table 2-4). This correction again assumes the same untested relationship between the number of channels per cell and the transcript ratio. The correction may also count the smaller size twice, because each transcript ratio is already relative to the RNA detected per nucleus, which in KI Purkinje nuclei was 0.79 of WT at 18 weeks (Methods). If so, the corrected conductances are too high. The recorded firing rate cannot show what, if anything, changed in KI Purkinje cells, because keeping Cav2.1 at its WT value and correcting for the smaller membrane each brought the firing rate into the recorded rate range. Different combinations of conductances can generate similar activity (Prinz et al. 2004; Goaillard and Marder 2021), so matching a specific readout does not identify the conductances.

What makes the interspike intervals of KI Purkinje cells more variable remains open. The models that we used omit calcium release from the endoplasmic reticulum. SK2 was left at its WT value because its transcript showed no significant change, but SK activity can fall without a change in transcript: in Cav2.1 mutant mice, the intervals become more variable as the calcium-activated potassium current falls, and a drug that opens SK channels reduces this variability (Walter et al. 2006). In KI Purkinje cells, a combination of chlorzoxazone, an activator of calcium-activated potassium channels, and baclofen, a GABA-B receptor agonist, lowered the CV at an intermediate dose without changing the firing rate (Bushart et al. 2021).

The models also omit the other changes of the SCA1 cerebellum. In KI Purkinje cells, the higher CV persisted with synaptic blockers and is therefore thought to arise within the Purkinje cell (Bushart et al. 2021), but the circuit around these cells also changes. In the KI line, basket cells make more inhibitory synapses on Purkinje cells, which at postnatal days 20–24 receive larger and more frequent inhibitory currents (Edamakanti et al. 2018), and Bergmann glia are reactive at 16 weeks (Edamakanti et al. 2023). In SCA1 mice, molecular layer interneurons become hyperactive early in disease (Pilotto et al. 2023), and in ATXN1[82Q] mice, the climbing fibre input to Purkinje cells is abnormal (Barnes et al. 2011). In our runs with synaptic input, the KI-scaled and WT simulated neurons received the same input, so these runs test the channel changes alone.

The simulations show that the firing rate could be brought into the recorded rate range by different assumptions about the Cav2.1 current and the membrane capacitance, so at least these two properties should be measured, the Cav2.1 current first. These measurements would replace two of the assumptions in the model with measured values.

### Limitations

The permutation p-values between mice are descriptive: with 3 mice per genotype and age, an exact two-sided permutation test cannot give a p-value below 0.1. Sex and age are confounded: the 18-week mice are male and the 30-week mice are female. This does not affect comparisons within an age, but may affect comparisons between ages. Since each mouse contributed only one library, we could not distinguish biological variance between mice from technical variance between libraries or test for any relationship between genotype and processing batch. When we projected the held-out mice onto the WT age trajectory (Methods), the 12-week WT mice projected to about 21–25 weeks, not between 5 and 18 weeks. Thus, the held-out mice may not be equivalent to the discovery mice, and we do not address the relationship of the KI state to normal ageing. Comparisons of genotypes at 12 weeks were made within the held-out mice. Nuclear RNA is enriched in newly transcribed RNA and lacks dendritic transcripts. Lastly, the genotype axes are based on a curated list of 1,414 cell–cell communication genes (not the whole transcriptome) and thus reflect only that part of the transcriptional state.

The Zang models were run at their published temperature of 34 °C (the Masoli model at 37 °C), and each simulated neuron was compared with its own WT run. Without synaptic input, only one noise level was used, at which the CV of the WT simulated neuron was lower than that of recorded WT Purkinje cells (0.02 vs 0.12 and 0.11). The population runs lasted 10 s, of which 5 s were scored, whereas the recorded CVs come from 150 s per cell (Stoyas et al. 2020), so the simulations can miss rare pauses. Each parameter set had one noise or synaptic seed, and the runs were scored by a single implementation. Every parameter set was scaled by the same transcript ratios, averaged over the nuclei and mice of each genotype, although KI Purkinje cells may differ in their channel losses. A model built from averaged conductances need not behave like the average cell (Golowasch et al. 2002). Block thresholds were determined from 2-s runs with a constant current, without the −80 mV holding potential used in the recordings of KI Purkinje cells. The block thresholds of KI Purkinje cells and the staining ratios were read from published figures. The gene-combination, staining-ratio, Cav2.1-step, membrane-area and block-threshold runs were designed after the result with all five transcript ratios was known, and are therefore reported descriptively only. The membrane-area correction changed only the conductance densities, not the morphology, and the capacitance that it relies on was measured at 12 weeks at the latest, whereas the KI Purkinje cells were recorded at 14 weeks. The transcript ratios are from 18 weeks, and the recordings of KI Purkinje cells and the staining from 14 weeks. At 12 weeks, in the held-out mice, the five transcripts were reduced by 17–32 %. The recorded phenotype comes from one study (Bushart et al. 2021) and differs between mouse lines (Dell’Orco et al. 2015), so any target for a model must specify the line and the age.

## Methods

### Data and cohort

All RNA-seq datasets used in this study are publicly available (Table 4).

**Table 4.** RNA-seq datasets.

| Dataset | Accession (reference) | Assay and tissue | Animals or donors | Use in this study |
|---|---|---|---|---|
| Discovery mice | GSE246183 (Tejwani et al. 2024) | snRNA-seq, cerebellum of Atxn1 154Q/2Q knock-in (KI) and wild-type (WT) mice | 24 mice: 3 KI and 3 WT at each of 5, 18, 24 and 30 weeks; 280,295 nuclei, 1,971 Purkinje | Discovery: every fit and every test |
| Held-out mice | GSE246183 (Tejwani et al. 2024) | snRNA-seq, cerebellum of Atxn1 154Q/2Q KI and WT mice | 6 mice: 3 KI and 3 WT at 12 weeks; about 38,000 nuclei | Replication in mice held out of the discovery analyses |
| ATXN1[82Q] transgenic | GSE215336 (Borgenheimer et al. 2022) | snRNA-seq, cerebellum of ATXN1[82Q] and WT mice | 3 mice per genotype at 12 weeks | Replication of the channel gene set in Purkinje nuclei |
| SCA7 knock-in | GSE269430 (Bartelt et al. 2024) | Purkinje-enriched snRNA-seq, cerebellum of SCA7 266Q and WT mice | 5 and 8.5 weeks; 4 mice per genotype and age | Exploratory replication in Purkinje nuclei |
| Knock-in bulk cerebellum | GSE122099 (Driessen et al. 2018) | Bulk RNA-seq, cerebellum of Atxn1 154Q/2Q and ATXN1[82Q] mice | 5 and 12 weeks; 3 mice per group | Replication in bulk tissue from the authors’ values; dilution test |
| ATXN1[82Q] bulk cerebellum | GSE75778 (Ingram et al. 2016) | Bulk RNA-seq, cerebellum of ATXN1[82Q] and WT mice | 5, 12 and 28 weeks; 3 mice per group | Secondary bulk comparison from the authors’ values |
| Human SCA1 | GSE246183 (Tejwani et al. 2024) | snRNA-seq, post-mortem cerebellum | 10 control and 10 SCA1 donors; 13 kept (7 control, 6 SCA1) with at least 10 Purkinje nuclei | Replication in human Purkinje nuclei |

### Criteria and reproducibility

Where an analysis or simulation was judged against a criterion, the criterion is given with the result and in the corresponding Methods subsection. The SCA7 analysis is called exploratory because it concerns a different disease gene and mouse model. We compared the simulations with the phenotype of KI Purkinje cells on four readouts: the firing rate ratio (recorded rate range, 0.86–1.33), the CV ratio, the afterhyperpolarization difference and the block-threshold ratio (Firing metrics and the reference phenotype). The gene-combination, staining-ratio, Cav2.1-step, membrane-area and block-threshold runs were designed after the result with all five transcript ratios was known, each in the light of the previous one, and are described without a pass or fail criterion. In the Masoli model, an increase in the CV of the interspike interval of a parameter set was counted only when it exceeded a bootstrap margin (see the subsection on the second published model). Each step was run once, and failed runs were kept on record with the reason for the failure. The code, its configuration files and the result tables and run records behind the figures and the text are released with the paper (Code accessibility).

### Differential expression

Counts were summed per mouse (pseudobulk) for each cell type and age. This gave three samples per genotype per age, and at each age KI was compared with WT using limma-voom, with the mouse as the unit. The main analysis comprised 27,415 tests (one per gene, cell type and age) across all cell types and ages, and hits were called at a Benjamini–Hochberg false discovery rate of 0.05. Gene-set tests were corrected separately from single-gene tests. The six 12-week mice were added in a separate five-age run after the four-age analysis had been completed and fixed. The four-age analysis is the main analysis, and the 12-week values are reported as replication. Gene sets, including the published SCA1 Magenta module and the channel gene set, were examined in the same framework. Per-age fits were used if a test of within-age variance suggested heterogeneity consistent with the sex-by-age design.

### Genotype axes and replication

Genotype axes were defined using the Purkinje nuclei of the 24 discovery mice in the Tejwani et al. (2024) data (3 KI and 3 WT at each of 5, 18, 24 and 30 weeks; 1,971 Purkinje nuclei in total). We fitted 30 principal components to depth-normalized log counts of a panel of 1,414 cell–cell communication genes (ligands, receptors, ion channels, pumps and transporters, and synaptic genes), of which 1,120 passed the expression filter in Purkinje nuclei and entered the fit. This panel was assembled before any comparison of KI with WT, from public ligand–receptor, channel and synapse resources (CellChatDB, NeuronChat, the Guide to Pharmacology, UniProt keywords and SynGO).

A principal component was considered a genotype axis if the three KI and three WT per-mouse means separated completely (same sign) at ≥3 of the 4 discovery ages. Complete separation means that all KI mice are on the same side of all WT mice. For each age, the axis direction was determined as the unit vector from the mean of the three WT per-mouse means to the mean of the three KI per-mouse means on the genotype axes that separated at that age, and nuclei were scored by their position along this direction for each age. A nucleus was defined as shifted if its score was above the 90th percentile of the scores of WT nuclei of the same age. The shifted fraction is the fraction of the nuclei of a mouse that meet this condition. By construction, it is about 0.10 in WT mice.

Replication was tested in the six held-out 12-week mice (203 Purkinje nuclei), whose nuclei were projected onto the fitted principal components without refitting. As no direction was fitted at 12 weeks, these nuclei were scored on the directions and thresholds of the two neighbouring discovery ages (5 and 18 weeks). The WT shifted fractions of the held-out mice (0.17–0.29) are above the 0.10 expected within an age because the threshold comes from WT nuclei of a different age. As a check on the held-out mice, we also projected their nuclei onto a WT age trajectory, the line joining the centres of the WT discovery mice at 5, 18, 24 and 30 weeks in principal-component spaces fitted to the WT discovery nuclei only (all detected genes, gene-set scores, and the genes enriched in Purkinje cells; 20 components each). In none of the three spaces did the 12-week WT mice fall between the 5-week and the 18-week WT mice (Limitations).

### Robustness of the Purkinje cell transcriptional state

Each mouse was excluded in turn. We kept the principal components but recomputed the axis direction and the WT 90th percentile from the other five mice of that age, and scored the nuclei of the left-out mouse on them. The shifted fraction of the left-out mouse was then calculated. The KI shifted fraction was 0.52, 0.95, 0.75 and 0.91 at 5, 18, 24 and 30 weeks, close to the values of the full fit (0.535, 0.950, 0.754 and 0.917), and the left-out WT mice scored 0.04–0.17 (mean 0.10).

In an effort to avoid learning the axis direction and evaluating the shifted fraction on the same counts, we split the UMI counts of each nucleus and gene separately into two halves by binomial sampling (p = 0.5; count splitting, Neufeld et al. 2024). The first half was used for fitting to obtain the axis direction and components, and the shifted fraction was measured on the second half. This was repeated for 10 random splits, and with negative-binomial splitting as a sensitivity test. The criterion was a KI shifted fraction of ≥0.50 at 18, 24 and 30 weeks. Averaged over the 10 splits, the KI shifted fraction was 0.49, 0.91, 0.79 and 0.88 at 5, 18, 24 and 30 weeks, which met the criterion, and the negative-binomial splits also passed the criterion.

We then split each component score into the part carried by the genes that Purkinje cells express at least as highly as any other cell type and the rest, and compared KI with WT on each part, to see if the difference between KI and WT was carried by the Purkinje cell’s own genes. The spread within a mouse is the standard deviation of the scores of the nuclei of that mouse on a component. For each of the 4 components, the slope of the spread across ages was greater in KI than in WT. To test whether the nuclei form two groups for each component, the scores of the nuclei of each mouse at 24 and 30 weeks were fitted with one- and two-component Gaussian mixtures. If at least two of the three mice of a genotype favoured the two-component fit at a component and age, we judged the genotype to have two groups.

### Replication in external datasets

We interrogated five publicly available datasets (Table 4): single-nucleus RNA-seq of Pcp2-ATXN1[82Q] and WT cerebellum at 12 weeks (GSE215336; Borgenheimer et al. 2022), Purkinje-enriched single-nucleus RNA-seq of SCA7 266Q and WT cerebellum at 5 and 8.5 weeks (GSE269430; Bartelt et al. 2024), single-nucleus RNA-seq of human SCA1 and control cerebellum (the human arm of GSE246183; Tejwani et al. 2024), and bulk RNA-seq of Atxn1 154Q/2Q and ATXN1[82Q] cerebellum (GSE122099, Driessen et al. 2018; GSE75778, Ingram et al. 2016). For the three single-nucleus datasets, we summed the counts of Purkinje nuclei per mouse or donor and compared KI or disease with control using limma-voom, as for the discovery data. For the two bulk datasets, we used the values reported by the authors and approximated their confidence intervals from the reported p values.

For the ATXN1[82Q] and human data, we identified Purkinje nuclei from the authors’ cell-type labels. Mice or donors with fewer than 10 Purkinje nuclei were excluded, which left 13 of the 20 human donors (29–76 Purkinje nuclei per mouse in the ATXN1[82Q] data; 749 Purkinje nuclei in total in the 13 donors). We identified Purkinje clusters in the SCA7 data from marker genes (250–626 Purkinje nuclei per mouse). Under the first rule, a cluster was labelled Purkinje if Pcp2, Calb1, Car8 and Itpr1 were all above their median across clusters and no marker of another cell type was above its median across clusters. In a Purkinje-enriched sample, this rule left no usable cluster, because the markers of other cell types were near zero almost everywhere. The second rule, which we used, kept the same positive markers and excluded a cluster if the mean log-normalized expression of Gabra6, Slc1a3, Gdf10, Kit, Mbp or Cx3cr1 was above 0.5. The deposited objects also carry the authors’ cell-type labels. All nuclei selected at 8.5 weeks are labelled Purkinje cells, and at 5 weeks all but two small clusters that the authors labelled endothelial (103 nuclei) and ependymal (27 nuclei), 4.6 % of the 2,796 selected nuclei (5.0 % in SCA7 and 4.3 % in WT mice). Because this share is small and similar in both genotypes, it can explain at most a small part of the 11–33 % reductions at 5 weeks. The second rule is unlikely to create the reduction by itself: of the nine test transcripts, it uses only that of IP3R1 (the marker gene Itpr1), and selecting nuclei with high Itpr1 in both genotypes makes the genotypes more similar for that gene, which works against finding a difference.

We also report, for each dataset, how many of the five channel transcripts and how many of the nine transcripts (the five channels plus IP3R1, SERCA3, BK β4 and α2δ2; genes Itpr1, Atp2a3, Kcnmb4 and Cacna2d2) were reduced in KI or disease (one-sided sign test over the nine). The sign test treats the nine transcripts as independent. Because they change together, its p values are nominal (0.002 in the ATXN1[82Q] single-nucleus data, in the SCA7 data at each age and in the ATXN1[82Q] bulk data at each age; 0.50 in the human data). In a sensitivity analysis including all 20 human donors (10 SCA1, 10 control), all five channel transcripts and all nine transcripts were lower (sign test over the nine, p = 0.002), but this analysis included donors with only 2–9 Purkinje nuclei, so we do not base our conclusion on it. We asked two further questions in the bulk data. To test whether the reduction is specific to the channel transcripts, we compared it with that of a control set of 113 Purkinje-enriched genes. To test dilution as the explanation for the bulk result, we computed, for each transcript, the change expected in bulk tissue under dilution (the fraction of its cerebellar transcripts that comes from Purkinje cells, estimated from the WT discovery mice, times its change in Purkinje nuclei at 12 weeks) and compared it with the observed bulk change by rank correlation. This correlation was 0.60 over the five channel transcripts, below the value of 0.8 that we had set as support for dilution.

### Firing metrics and the reference phenotype

Somatic spikes were detected as upward crossings of 0 mV in the Masoli model and of −30 mV at the soma in the Zang models, with a refractory period of 2 ms. Dendritic calcium spikes were detected as upward crossings of −30 mV at a distal dendritic site, with a refractory period of 20 ms. The somatic voltage of a simulated neuron is the membrane potential at the soma, as measured in a whole-cell current-clamp recording. The firing rate and the CV of the interspike interval were computed from the spike times only, as in the cell-attached recordings of the reference phenotype. The afterhyperpolarization and the block threshold were taken from the somatic voltage, as in its whole-cell recordings.

From the spike times in the scoring window, we computed the firing rate (the number of spikes divided by the length of the scoring window), the coefficient of variation (CV) of the interspike interval (ISI; its standard deviation divided by its mean) and CV2 (the mean of 2|ISI(i + 1) − ISI(i)| / (ISI(i + 1) + ISI(i)) over consecutive intervals). A pause is defined as an interval longer than three times the median interval of the run. The afterhyperpolarization is the lowest somatic voltage between two consecutive spikes, taken as the median over the intervals that are not pauses. The scoring window was 5–10 s for the 10-s population runs, 10–25 s for the 25-s runs and 30–45 s for the Masoli runs.

A 10-s population run was scored as in depolarization block if it had fewer than three spikes in the scoring window and a mean somatic voltage above −55 mV. With fewer than three spikes and a mean voltage at or below −55 mV, it was scored as at rest. In the block-threshold series (2-s runs with a constant injected current), a run was scored as in block if there was no spike in the last 0.25 s and the mean somatic voltage over that time was above −55 mV. The block threshold is the lowest injected current at which a run is in block. In this series, the afterhyperpolarization of a KI-scaled run was compared with that of WT at the same firing rate, with the WT value interpolated between the WT runs on either side of that rate.

The reference phenotype is that of Bushart et al. (2021): Purkinje cells of Atxn1 154Q/2Q mice at 14 weeks, recorded in acute slices at 33 °C, in lobules IV and V (anterior cerebellum) and in lobule X (nodular zone). The values are group means from Supplementary Table 1 of that paper: firing rates of 31.65 (WT) and 42.04 (KI) spikes per second in the anterior cerebellum and of 27.81 and 23.78 in the nodular zone (KI/WT 1.33 and 0.86, neither significant); a CV of the interspike interval of 0.1171 and 0.1145 in WT and of 0.2082 and 0.3274 in KI (KI/WT 1.8 and 2.9); an afterhyperpolarization of −65.93 mV (WT) and −63.88 mV (KI) in the anterior cerebellum and of −66.65 and −65.03 mV in the nodular zone, that is, 2.05 and 1.62 mV less negative in KI; and no significant difference in input resistance (WT 53.5 MΩ and KI 57.2 MΩ in the anterior cerebellum; WT 74.0 MΩ and KI 80.0 MΩ in the nodular zone). The same table shows no significant difference between KI and WT Purkinje cells in spike peak, rates of rise and fall, or spike half-width. We did not compare the spike waveform with the simulated neurons. The block thresholds (1,260 pA in WT and 853 pA in KI in the anterior cerebellum, 1,654 and 840 pA in the nodular zone, that is, 0.68 and 0.51 of WT) and the staining ratios (BK 0.79 and 0.77, Cav3.1 0.77 and 0.73, Cav2.1 1.01 and 0.93, and the IP3 receptor 0.97 and 0.98 of WT in the anterior cerebellum and nodular zone; BK and Cav3.1 significantly reduced, Cav2.1 and the IP3 receptor not; Kv3.3 and Kv4.3 not measured) were read from the published figures. Recording conditions were those of Stoyas et al. (2020): firing rate and CV were measured in the cell-attached configuration over 150 s, and the afterhyperpolarization in whole-cell mode, as the most negative voltage after each spontaneous spike. For the block threshold, the cells were held at −80 mV and injected with depolarizing current.

For each KI-scaled run, the WT run of the same parameter set and seed served as the baseline for four readouts: the firing rate ratio (KI/WT), the CV ratio, the afterhyperpolarization difference (KI − WT; positive if less negative) and the block-threshold ratio. Pauses and runs in depolarization block were counted and reported.

### Purkinje cell model and simulations

We used the model of Zang et al. (2018) (ModelDB accession 243446) and simulated it in NEURON 8.2.6 at 34 °C, on one thread. The model has 1,137 sections. The time step was 0.02 ms, and no holding current was applied. Without synaptic input, the interspike intervals of the model are almost constant, so a noise current was injected at the soma (an Ornstein–Uhlenbeck process with a standard deviation of 6.5 pA and a time constant of 5 ms), with one noise seed per parameter set.

In every section that carries a given mechanism, the conductance of that mechanism was scaled by the matching transcript ratio: Kv3.3 (Kcnc3) scaled Kv3; Kv4.3 (Kcnd3), Kv4 and Kv4s; BK (Kcnma1), mslo and abBK; Cav2.1 (Cacna1a), newCaP; and Cav3.1 (Cacna1g), CaT3_1. The scaling factor was the KI/WT transcript ratio at 18 weeks (the age of the discovery mice closest to that of the KI Purkinje cell recordings, 14 weeks): Kv3.3 0.667, BK 0.719, Kv4.3 0.588, Cav2.1 0.672 and Cav3.1 0.649.

To vary the model, we drew 200 parameter sets, scaling each of eight conductance groups (resurgent sodium, Cav2.1 (P/Q-type) calcium, Cav3.1 (T-type) calcium, Kv3, Kv4, BK, SK2 and HCN) by a factor between 1/1.5 and 1.5 (Latin hypercube on a logarithmic scale, with a fixed random seed). Each set was run for 10 s and scored on 5–10 s. A set was kept if its WT run fired at 20–120 Hz with a CV of the interspike interval below 0.2 and without a plateau (a stretch longer than 1 second above −30 mV). This left 130 parameter sets, and every KI-scaled run was paired with the WT run of the same parameter set and seed.

For runs with synaptic input, we used the model of Zang and De Schutter (2021; ModelDB accession 266864) as published, at 34 °C: 2,000 parallel-fibre synapses firing at 0.135 Hz and 144 stellate-cell and 160 basket-cell synapses firing at 14.4 Hz, with no climbing-fibre input, no noise current and no holding current. The 2021 model is not simply the 2018 model with synapses added: in the spiny dendrites its Kv3 conductance is 0.667 of the 2018 value, its mslo conductance is twice as large and abBK is added, and in the main dendrites its HCN conductance is 1.667 times that of the 2018 model. We ran the same 130 parameter sets, with their eight scale factors applied to the published values of the 2021 model, in 10-s runs scored on 5–10 s, with one synaptic seed per parameter set. Each KI-scaled run was paired with the WT run of the same parameter set and seed, and the published parameter set was run separately.

On the same 130 parameter sets and seeds, we ran, without synaptic input, all 32 combinations in which each of the five conductances was either scaled by its 18-week transcript ratio or left unchanged; the staining ratios (BK 0.78 and Cav3.1 0.75 of WT, averaged over the two regions; Bushart et al. 2021), alone and with the Kv3.3 or Kv4.3 transcript ratio or both added (Extended Data Figure 6-3); the staining ratios with the Cav2.1 conductance lowered in steps to 0.95, 0.90, 0.85 and 0.80 (Extended Data Figure 6-2); and six membrane-area versions (A = 0.9 to 0.6). With synaptic input, we ran the versions reported in the Results, each conductance reduced alone and each restored alone. Every run is listed in Extended Data Tables 2-2 and 2-3. The runs without synaptic input are the main comparison, because the higher CV of KI Purkinje cells persisted in the presence of synaptic blockers (Bushart et al. 2021). Figure 6B shows 35 versions without synaptic input (the 31 combinations other than WT and the four staining-ratio versions) and 14 with synaptic input (all five transcript ratios, the five versions with one conductance unchanged, the five with one conductance reduced alone, and three staining-ratio versions).

To test whether the effect of a reduction depends on the other conductances, we compared, for each conductance, every pair of versions that differ only in that conductance: 16 pairs among the 32 combinations, and 18 for Kv3.3 and for Kv4.3, which were also varied in the staining-ratio versions. Each pair was compared in each of the 130 parameter sets, giving at most 16 × 130 = 2,080 comparisons (18 × 130 = 2,340 for Kv3.3 and Kv4.3). A comparison was discarded when the simulated neuron did not fire in one of the two runs, which left 1,979 comparisons for Cav2.1, 2,277 for Kv4.3, 2,009 for BK, 2,247 for Kv3.3 and 2,022 for Cav3.1. The Cav2.1 reduction raised the firing rate in 98.9 % of its comparisons, the Kv4.3 reduction in 98.6 % and the BK reduction in 95.9 %, whereas the Kv3.3 and Cav3.1 reductions lowered it in 98.2 % and 98.6 %. To test whether the reductions combine independently, we divided, for each of the 26 versions with two to five reductions and for each parameter set, the observed firing rate ratio by the product of the firing rate ratios obtained with each of its reductions alone. A quotient of 1 means exact agreement. The median quotient was 0.98–1.07 across these versions, and 1.05 with all five reductions. For the afterhyperpolarization, the change of a combined version was compared with the sum of the changes of its single reductions. It differed from that sum by −0.07 to +0.14 mV (Extended Data Table 2-1). Reduced alone, the afterhyperpolarization became less negative mainly with the Cav2.1 reduction and, to a smaller degree, with the BK reduction. Each of the other three reductions changed it by less than 0.5 mV.

The threshold to depolarization block was measured with a constant somatic current, switched on at the start of a 2-s run, together with the noise current described above, with one run per current and version. On the published parameter set, the current went from 0.25 to 3.0 nA in steps of 0.125 nA. On each of the 130 parameter sets, ten currents were used (0.25 to 2.0 nA in steps of 0.25 nA, then 2.5 and 3.0 nA), for WT, the version with all five transcript ratios, the version with Cav2.1 unchanged, the staining-ratio versions and the membrane-area versions.

A preliminary test in a two-compartment model (Forrest 2015; ModelDB accession 180789) was set aside, because it targeted a depolarized plateau, which is not the firing recorded in Purkinje cells of the KI line (Bushart et al. 2021), and because that model has no named mechanism for several of the genes of interest.

### Second published model (Masoli et al. 2015)

We used the model of Masoli et al. (2015) (ModelDB accession 229585), which has a complete morphology and a named mechanism for each of the five channels, at its published temperature of 37 °C. The conductances of the five channel mechanisms (Kv3.3 and Kv4.3 in the dendrites, BK, Cav2.1 and Cav3.1) were multiplied by their KI/WT transcript ratios, using the point estimate and the two ends of its 95 % confidence interval. We drew 40 WT variants by scaling the conductances of eight channel types (the five channels, Nav1.6, HCN1 and SK2) log-uniformly between 0.5× and 1.5× their published values. Of these, 14 fired at 20–120 Hz with an ISI CV below 0.2 (25 were silent and 1 fired at 122 Hz), and we continued with these 14. A somatic noise current (an Ornstein–Uhlenbeck process with a time constant of 5 ms and σ = 6.5 pA) made the interspike intervals variable, and each KI-scaled run used the noise seed of its own WT run. Runs lasted 45 s and were scored on 30–45 s. Because the simulated neurons fire at 46–117 Hz at 37 °C, each KI-scaled run was compared with its own WT run rather than with recorded values. We counted an increase in the ISI CV when the ISI CV of the KI-scaled run exceeded that of its own WT run by more than twice their combined bootstrap standard error (moving-block bootstrap of the interspike intervals, blocks of 10 intervals, 2,000 resamples), and never by less than the rounding error of spike times recorded on a 0.1 ms grid.

With the 18-, 24- and 30-week ratios, the median firing rate ratio was 1.14, 1.11 and 1.12, no parameter set had an increased ISI CV (0 of 14) and no run produced a pause. With the 5-week ratios, with or without the HCN1 increase of that age, the firing rate changed by less than 2 %, and no parameter set had an increased ISI CV. Restoring Cav2.1 alone lowered the firing rate ratio in 14 of the 14 parameter sets (Extended Data Figure 6-1). KI/WT ratios taken from the most shifted KI nuclei and exaggerated ratios gave no more variable interspike intervals and no pauses (Extended Data Figure 6-4). An independent implementation reproduced the firing rates, CV2 values and pauses of these runs from the spike times, except those of the runs with exaggerated ratios (Independent re-derivation), and the ISI CV values come from a single implementation.

### Membrane-area correction and α2δ2

For the membrane-area correction, each of the five conductance factors was set to r/A, where r is the 18-week KI/WT transcript ratio of the channel and A is the membrane area of the KI cell as a fraction of WT. All other conductances and the morphology were unchanged. A was varied from 0.9 to 0.6. In the KI line, the capacitance of Purkinje cells has been reported at 6–11 weeks (0.77 of WT for the whole cell and 0.75 for the dendritic tree, calculated from Table 1 of Watase et al. 2002) and at 12 weeks (534.1 ± 27.6 pF in 22 KI cells from 8 mice, against 670.4 ± 26.7 pF in 20 WT cells from 8 mice, that is 0.80 of WT; Table 1 of Shuvaev et al. 2022), but not at 14 weeks, the age of the KI Purkinje cell recordings. This correction assumes that the number of channels of each of the five kinds per cell follows its transcript ratio, that the densities of all other channels are unchanged, and that the RNA content of the nucleus is unchanged. In our data, the RNA content of KI Purkinje nuclei (the median UMI count per nucleus, relative to that of granule cell nuclei of the same mouse) was 0.79 of WT at 18 weeks.

The reduction of α2δ2 was represented as a factor of 0.85 on the Cav2.1 conductance (the newCaP mechanism). This is the 35 % lower calcium-channel current density (barium current) of Purkinje cells dissociated from 4- to 8-day-old ducky mice, which lack full-length α2δ2 (Barclay et al. 2001), scaled by the fraction of the α2δ2 transcript (Cacna2d2) lost in KI Purkinje nuclei (0.42). This scaling assumes that the current falls in proportion to the transcript. In heterozygous ducky mice, the current density is not reduced (Barclay et al. 2001), so 0.85 is a value for a sensitivity test, neither an independent estimate nor a bound on the loss in KI Purkinje cells.

### Independent re-derivation

An independent implementation, written from the stated criteria without access to the code or results of the first implementation, recomputed the following from the raw inputs, and the two were compared at the precision reported: the principal components and the shifted fractions, with leave-one-mouse-out and two of the count splits of each kind; the RNA content of the nuclei; and the bulk replication with its control set and the dilution test. The population runs (Table 2, Extended Data Tables 2-1 and 2-4, and Extended Data Figures 6-2 and 6-3) were scored by a single implementation. The firing rates, the CV2 values and the pauses of the Masoli runs (Extended Data Figure 6-4) and the values of the five-seed runs of the published parameter set of the Zang et al. (2018) model (Extended Data Figure 5-1) were reproduced from the spike times in the same way, except those of the Masoli runs with exaggerated ratios. The ISI CV of the Masoli runs and the count of parameter sets with an increased ISI CV were computed by a single implementation. Three limma comparisons were re-implemented after the values of the first implementation had been seen, following a correction to the specification. They are therefore reported as independent re-implementations, not as blind checks. The 12-week limma comparison has not been independently recomputed.

Use of AI tools. Omkar Chandra wrote the manuscript. The analysis and simulation code, including the independent re-implementations, was written with AI coding tools (Claude, Anthropic; ChatGPT, OpenAI; GLM, Z.ai), directed by him. He specified every analysis and its criteria and verified all code, inputs and reported values. The same tools were used to draft parts of the Methods and figure legends and to edit the text for language. All text was revised and approved by the authors.

### Statistics

Between-mouse comparisons used exact permutation tests where reported. With three mice per genotype and age, the smallest two-sided p-value is 0.1, so these p-values are descriptive. Counts of components, measures and models meeting a stated rule are reported against their chance expectation where one exists (Criteria and reproducibility). Differential expression used limma-voom in R (R 4.4.3–4.5.3, limma 3.62.1–3.66.0, edgeR 4.4.0–4.8.2 across runs). Single-nucleus quality control and the state analyses were written in Python with anndata, numpy, pandas, scipy and h5py (the quality-control run recorded scanpy 1.12.4, anndata 0.13.3.post0, numpy 2.5.3, pandas 3.0.5, scipy 1.18.1 and h5py 3.16.0), and the later Purkinje analyses used Python 3.14.4 with numpy 2.5.3, pandas 3.0.5, scipy 1.18.1 and h5py 3.16.0. The WT age trajectory was computed with Python 3.9.23 and scanpy 1.10.3, with gene-set scores from UCell 2.6.2 in R 4.3.1. Simulations used NEURON 8.2.6 under Python 3.10.1 with numpy 1.26.4. The versions recorded for each script are listed in the README of the code (Code accessibility).

## Code accessibility

The code described in the paper is provided as Extended Data 1 and at https://github.com/omkarchandra/sca1-transcript-conductance under the MIT License, and will be archived with a DOI on publication. The software and its versions are given under Statistics. The simulations were run on a Linux high-performance computing cluster with the Slurm scheduler.

## Data and code availability

The snRNA-seq data are available from GEO (GSE246183; Tejwani et al. 2024), the external datasets from GEO (GSE122099, GSE75778, GSE215336 and GSE269430). The Purkinje cell models are available from ModelDB (229585, Masoli et al. 2015; 243446, Zang et al. 2018; 266864, Zang and De Schutter 2021; 180789, Forrest 2015). The analysis and simulation code, its configuration files, the figure scripts and the result tables and run records behind the figures and the text are in the GitHub repository given under Code accessibility, which will be archived with a DOI on publication.

Author contributions: Omkar Chandra designed the study, wrote the code, performed the analyses and simulations, interpreted the results and wrote the manuscript. Caleb M. Smith contributed to the study design and reviewed the manuscript. Abbreviated title: SCA1 channel transcripts and Purkinje cell firing. Conflict of interest: The authors declare no competing financial interests.

## Extended Data

Extended Data Figures 2-1, 2-3, 3-2, 5-2 and 6-1 are in the extended data document. Extended Data Figures 2-2, 3-1, 4-1, 5-1, 6-2, 6-3 and 6-4 and Extended Data Tables 2-1, 2-2, 2-3 and 2-4 are sheets of one spreadsheet file, which also lists the thresholds of Extended Data Figure 6-1A. Each item has its full legend with it.

Extended Data Figure 2-1. Validation in the held-out 12-week mice.

Extended Data Figure 2-2. Gene loadings of the four genotype components of Purkinje nuclei, the mean score of each mouse on the four components, and the KI − WT difference of each component at each age.

Extended Data Figure 2-3. Distribution of Purkinje nuclei along the axis direction in each mouse.

Extended Data Figure 3-1. Differential expression of the channel and calcium-store genes of Figure 3 in Purkinje nuclei at five ages, including 12 weeks, and the gene-set test of the published SCA1 Magenta module (Ingram et al. 2016).

Extended Data Figure 3-2. Staining ratio against transcript ratio for the four proteins stained in KI mice.

Extended Data Figure 4-1. Replication: log2 fold change, 95 % confidence interval and q value of the nine transcripts in each comparison of Figure 4.

Extended Data Figure 5-1. Zang et al. (2018) model without synaptic input, published parameter set: every run of five noise seeds in eight conditions.

Extended Data Figure 5-2. Somatic voltage of the published parameter set with synaptic input, in WT and five versions.

Extended Data Table 2-1. All 32 combinations of the five transcript ratios on the 130 parameter sets, without synaptic input. Each row is one version: the conductances named in the first column were scaled by their 18-week KI/WT transcript ratios, and the others were left unchanged. Firing rate ratio: KI over WT, quartiles and median over the parameter sets that fire in both runs. In range: number of the 130 parameter sets with a firing rate ratio within the recorded rate range (0.86–1.33). CV ratio and AHP change: medians. The last three columns give the number of parameter sets in which the KI-scaled run fires, fires with pauses, or is in depolarization block.

Extended Data Table 2-2. Zang and De Schutter (2021) model with background synaptic input: every run of the 130 parameter sets and of the published parameter set in 21 versions.

Extended Data Table 2-3. Zang et al. (2018) model without synaptic input: the eight scale factors of the 200 drawn parameter sets, and every run of the 130 kept parameter sets in 46 versions.

Extended Data Table 2-4. Membrane-area correction of the five transcript ratios. Extended Data Figure 6-1. KI/WT transcript ratios in a population of Masoli models.

Extended Data Figure 6-2. Stepwise reduction of the Cav2.1 conductance on top of the staining ratios.

Extended Data Figure 6-3. Kv3.3 and Kv4.3 transcript reductions added to the staining ratios.

Extended Data Figure 6-4. Masoli model population: the runs of the 14 parameter sets in 23 conditions, the runs with the KI/WT ratios taken from the most shifted KI nuclei, and the runs with exaggerated KI/WT ratios.

Extended Data 1. Code (ZIP file): the analysis and simulation code, small frozen inputs and result tables, with a README that lists the scripts behind each figure and table and the software versions recorded for them.

## Notes

https://github.com/omkarchandra/sca1-transcript-conductance

### Competing Interest Statement

The authors have declared no competing interest.

