## Extended Data document (Figures 2-1, 2-3, 3-2, 5-2 and 6-1) for "Ion-channel transcript loss and variable firing in SCA1 Purkinje cells: a modelling test"

#### Extended Data in this document

##### Extended Data Figure 2-1

| component | *discovery KI − WT sign by age (weeks)* 5 | 18 | 24 | 30 | predicted | 12 weeks KI − WT | sign match | complete separation |
|---|---|---|---|---|---|---|---|---|
| component 5 | + | + | + | + | + | +2.49 | ✓ | ✓ |
| component 6 | − | − | − | − | − | −0.761 | ✓ | ✓ |
| component 11 | + | + | + | − | + | +0.296 | ✓ | ✗ |
| component 30 | − | − | − | − | − | −0.233 | ✓ | ✓ |

Extended Data Figure 2-1. Validation in the held-out 12-week mice. For each of the 4 Purkinje genotype components: the KI−WT sign in the discovery mice at 5, 18, 24 and 30 weeks; the predicted sign (the sign at most of the 4 ages); the 12-week KI−WT difference (mean of the 3 KI minus mean of the 3 WT per-mouse means); whether the sign matched; and whether KI and WT separated completely. The signs matched for all 4 components. With a one-half chance of each sign matching, we would expect about 2 of the 4 to match if genotype had no effect. KI and WT separated completely in the predicted direction for 3 of the 4 components. There are 20 ways for 6 mice to be split into 2 groups of 3, and only 1 of them is complete separation in the predicted direction, so by chance about 0.2 of the 4 components would be expected to separate completely in the predicted direction.

### Extended Data Figure 2-3

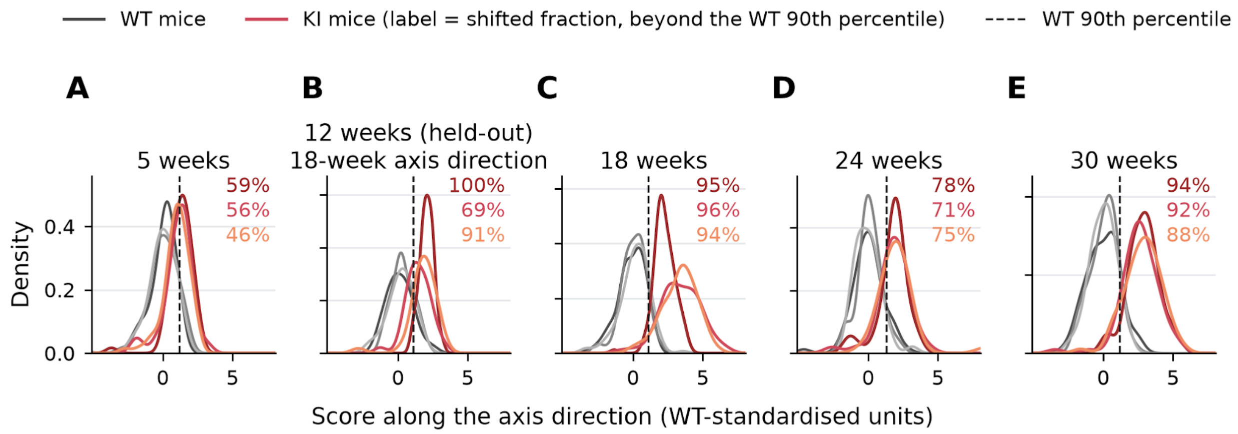


Extended Data Figure 2-3. Distribution of Purkinje nuclei along the axis direction in each mouse. For each age, kernel density of the nuclei's scores along the WT-to-KI axis direction (full-data fit; WT-standardized units), one line per mouse: WT, grey; KI, coloured, with each KI mouse's shifted fraction (share of its nuclei beyond the WT 90th percentile, dashed line) printed in its colour. (A) 5 weeks; (B) the six held-out 12-week mice, scored without refitting on the 18-week direction; (C) 18, (D) 24 and (E) 30 weeks. From 18 weeks, the distribution of each KI mouse is shifted away from that of WT, without a clearly separate WT-like peak. n = 3 mice per genotype and age; 2,174 nuclei (1,971 discovery, 203 held-out).

### Extended Data Figure 3-2

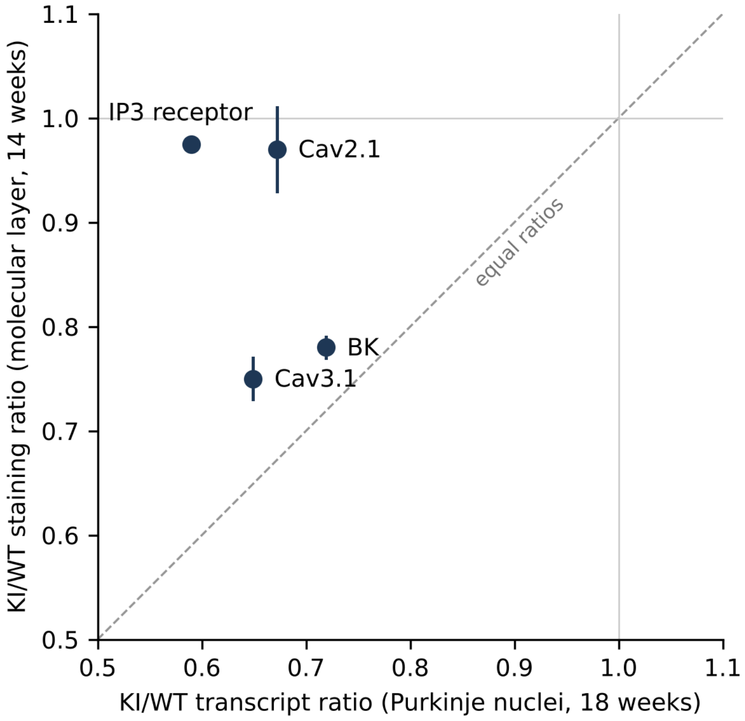


Extended Data Figure 3-2. Staining ratio against transcript ratio for the four proteins stained in KI mice. KI/WT staining ratio of the molecular layer at 14 weeks (Bushart et al. 2021; values read from their figure) against the KI/WT transcript ratio in Purkinje nuclei at 18 weeks (Figure 3), for BK, Cav3.1, Cav2.1 and the IP3 receptor (IP3R1 transcript). Point, mean of the anterior cerebellum and the nodular zone; vertical bar, the two regions; dashed line, equal ratios. All four transcripts were reduced (ratios 0.59–0.72). BK and Cav3.1 staining was also reduced (0.79 and 0.77 for BK, 0.77 and 0.73 for Cav3.1), whereas Cav2.1 and IP3 receptor staining was not (1.01 and 0.93 for Cav2.1, 0.97 and 0.98 for the IP3 receptor). Across the four proteins, the staining ratio did not rise with the transcript ratio, and four proteins are too few to estimate a correlation.

### Extended Data Figure 5-2

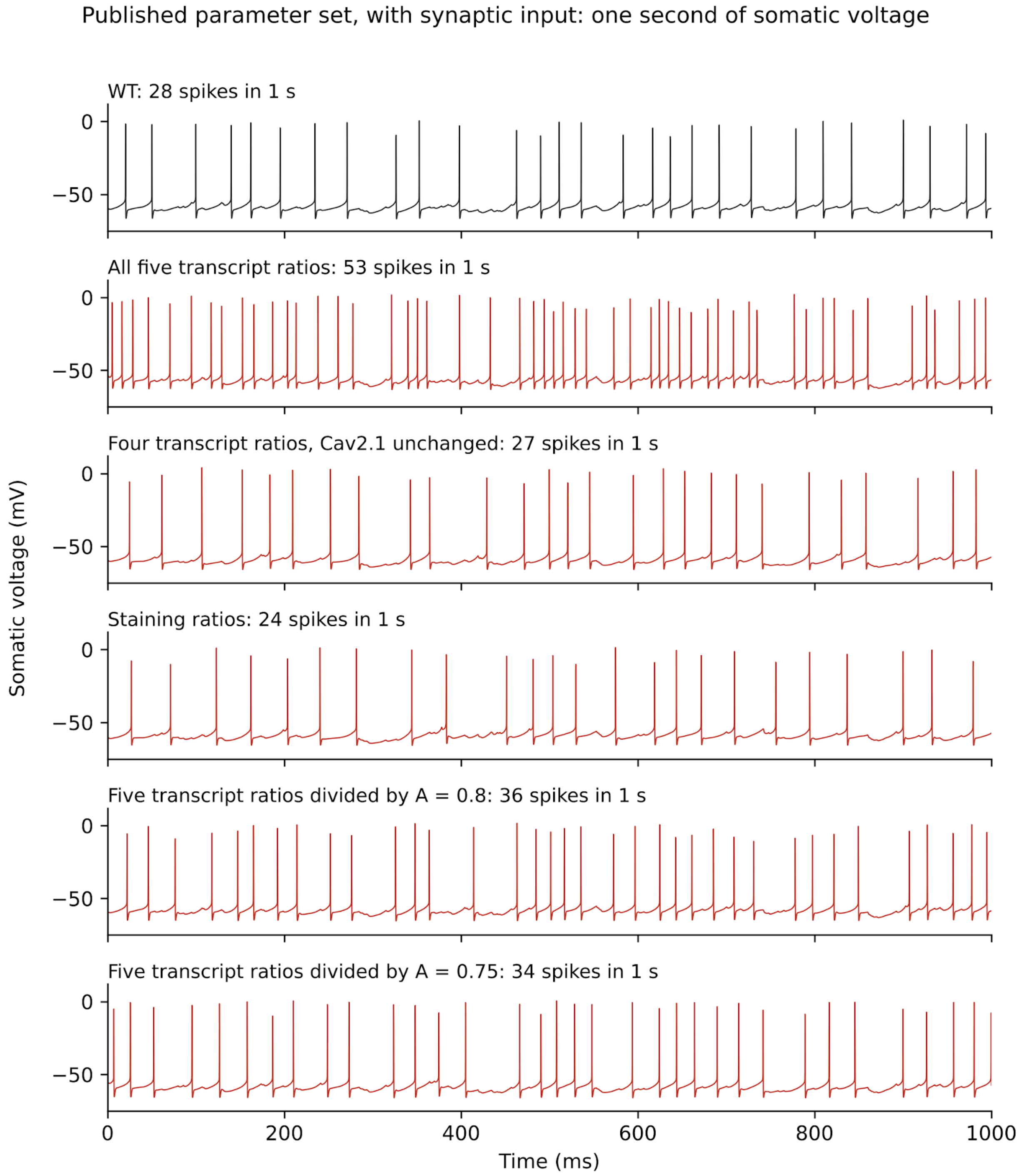


Extended Data Figure 5-2. Somatic voltage of the published parameter set with synaptic input, in WT and five versions. The traces are the membrane voltage at the soma, as in a whole-cell current-clamp recording. The firing rate and CV recorded in KI Purkinje cells (Bushart et al.

2021) were measured cell-attached, from spike times only. One second (5 to 6 s of a 10-s run) of the somatic voltage of the published parameter set of the Zang and De Schutter (2021) model with background synaptic input, in WT and in the five versions of Figure 5B: all five transcript ratios; the four transcript ratios of Kv3.3, BK, Kv4.3 and Cav3.1 with Cav2.1 unchanged; the staining ratios (BK 0.78 and Cav3.1 0.75 of WT, with Kv3.3, Kv4.3 and Cav2.1 at their WT values); and the five transcript ratios divided by the membrane area A = 0.8 and 0.75. Black, WT; red, KI-scaled. Each title gives the number of spikes in the second shown (upward crossings of −30 mV, as in the Methods). The WT and all-five traces are those of Figure 5A, bottom row. The traces show one parameter set; values over the 130 parameter sets are in Figure 5B and Table 2.

### Extended Data Figure 6-1

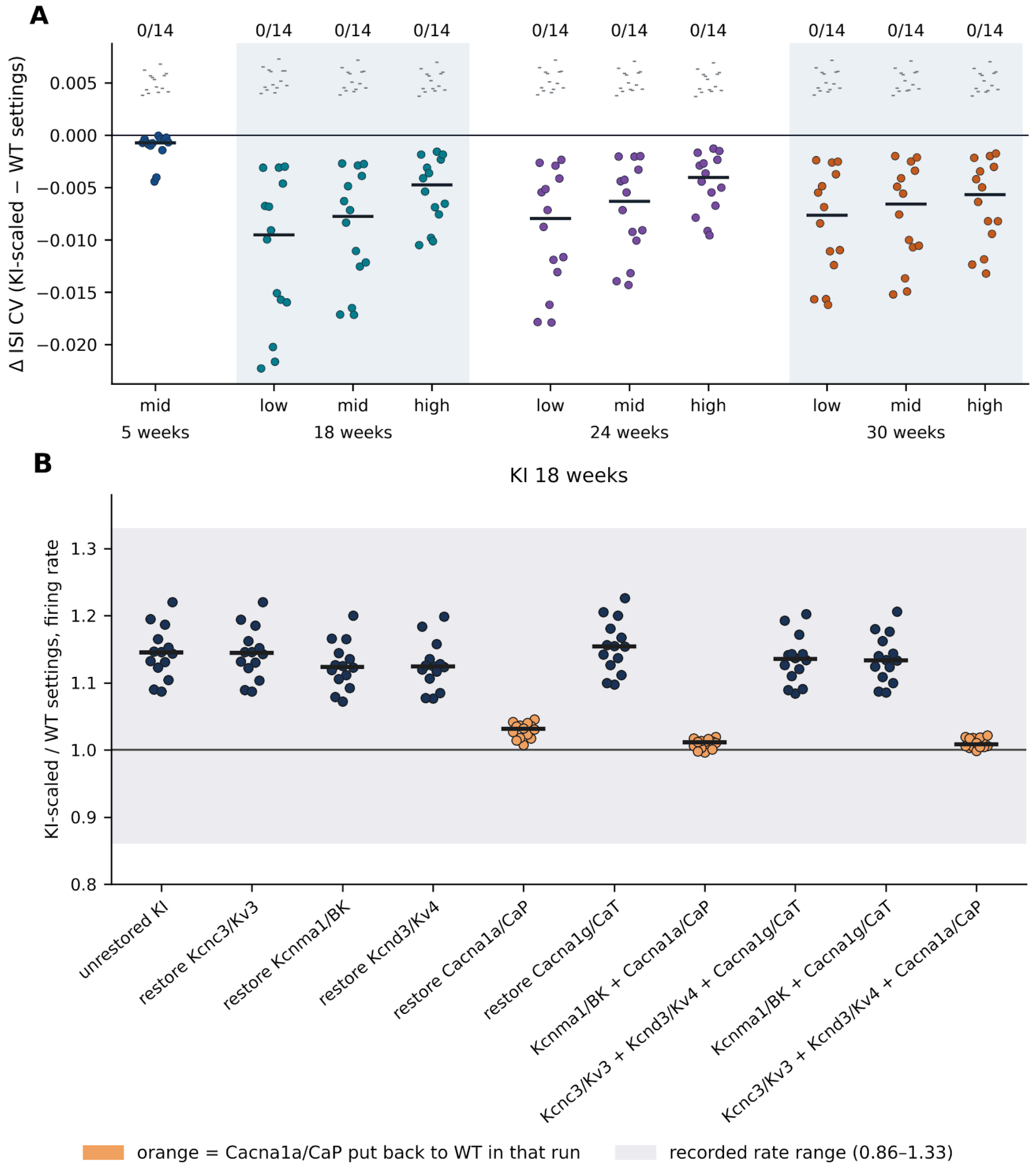


Extended Data Figure 6-1. KI/WT transcript ratios in a population of Masoli models. (A) Change in the CV of the interspike interval (KI-scaled minus own WT). Each dot is one of the 14 parameter sets. Columns are the low, mid and high estimates of the ratios (the point estimate and the two ends of its 95 % confidence interval) at 18, 24 and 30 weeks, and the mid estimate at 5 weeks; black bar, median. The small grey ticks above each column mark, for each parameter set, the threshold for counting an increase in the ISI CV: twice the combined bootstrap standard error of the two runs, or the rounding floor of spike times recorded on a 0.1 ms grid where that is larger (the floor set 119 of the 140 thresholds, all listed in the spreadsheet, sheet Figure 6-1A thresholds). The number above each column gives how many of the 14 parameter sets exceeded it (0 of 14 everywhere). (B) KI at 18 weeks: KI/WT firing rate ratio for the unrestored KI and for each single and grouped restoration of a channel to its WT value; orange, runs in which Cav2.1 was restored; shaded band, the recorded rate range (0.86–1.33). The values are in Extended Data Figure 6-4.

#### Extended Data provided as a spreadsheet

Extended Data Figures 2-2, 3-1, 4-1, 5-1, 6-2, 6-3 and 6-4 and Extended Data Tables 2-1, 2-2, 2-3 and 2-4 are provided as one spreadsheet file, sca1_transcript_conductance_supplementary_tables_v6_biorxiv, which has a Contents sheet.
